# The energetic stress cytokine GDF15 is elevated in the context of chronic and acute psychosocial stress

**DOI:** 10.1101/2024.04.19.590241

**Authors:** Qiuhan Huang, Anna S. Monzel, Shannon Rausser, Rachel Haahr, Claire E. Indik, Micah J Savin, Natalia Bobba-Alves, Cynthia C. Liu, Jack Devine, Elizabeth Thompson, Catherine Kelly, Mangesh Kurade, Jeremy Michelson, Shufang Li, Kris Engelstad, Vincenzo Lauriola, Anna L Marsland, Brett A Kaufman, Marie-Pierre St-Onge, Richard Sloan, Robert-Paul Juster, Gilles Gouspillou, Michio Hirano, Martin Picard, Daniel W. Belsky, Caroline Trumpff

**Affiliations:** Division of Behavioral Medicine, Department of Psychiatry, Columbia University Irving Medical Center, New York, NY, USA; Robert N Butler Columbia Aging Center and Department of Epidemiology, Columbia University Mailman School of Public Health, New York, NY, USA; Department of Neurology, H. Houston Merritt Center, Columbia Translational Neuroscience Initiative, Columbia University Medical Center, New York, NY, USA; Department of Psychology, University of Pittsburgh, Pittsburgh, PA, United States; Department of Medicine, Division of Cardiology, Center for Metabolism and Mitochondrial Medicine, University of Pittsburgh School of Medicine, Pittsburgh, PA United States; Division of General Medicine and Center of Excellence for Sleep & Circadian Research, Department of Medicine, Columbia University Irving Medical Center, New York, USA; Department of Psychiatry and Addiction, University of Montreal, Montreal, QC, Canada; Département des sciences de l’activité physique, Faculté des sciences, University of Quebec at Montréal (UQAM), Montréal, Quebec, Canada; New York State Psychiatric Institute, New York, NY, USA

**Keywords:** psychological stress, psychobiology, mitochondria, GDF15, DNA methylation, adversity, SES, stress physiology, epigenetic age

## Abstract

Growth Differentiation Factor 15 (GDF15) is a protein that reflects mitochondrial energetic stress and is linked to physical and mental health symptoms, aging, and mortality. Here, we tested the hypothesis that GDF15 is a stress-responsive biomarker through a series of observational and experimental studies. We report four main findings. First, in the UK Biobank (n=53,026) and Framingham Heart Study (FHS) Offspring (n=3,460) cohorts, plasma GDF15 levels were elevated in individuals with symptoms of depression and anxiety. In the FHS cohort, GDF15 was also higher in participants exposed to chronic psychosocial stressors, including lower educational attainment, lower family income, and higher job strain. Second, plasma GDF15 levels in the FHS cohort correlated positively with epigenetic clocks measuring biological aging and effect sizes of GDF15 associations with psychosocial stressors were comparable to those observed for the clocks. Third, in a two-participant intensive-sampling study (n=112 days), saliva GDF15 showed a robust awakening response similar to established stress-related hormones. However, it exhibited a distinct negative pattern, peaking at waking and declining by 42–92% within 30–45 minutes. Finally, in two laboratory experiments (n=148), acute social-evaluative stress significant increased GDF15 levels in plasma and saliva within minutes. Together, these findings suggest that psychosocial stress may contribute to mitochondrial energetic stress indexed by GDF15, with implications for aging and health. This work opens new avenues for using GDF15 as a non-invasive biomarker to study the biological embedding of stress and its impact on aging trajectories.

**Significance statement:** Growth Differentiation Factor 15 (GDF15) is a circulating protein elevated with mitochondrial energetic stress, aging, and diseases. Our findings show that GDF15 is elevated with depressive and anxiety symptoms and in those exposed to chronic psychosocial stress. Elevated plasma GDF15 also correlates with accelerated biological aging, as measured by epigenetic clocks. The effect sizes linking GDF15 to psychosocial stressors were comparable to those observed for the epigenetic clocks. Saliva GDF15 shows a robust negative awakening response characterized by elevated levels at awakening before declining within 30–45 minutes. Acute social-evaluative stress induced increase in plasma and saliva GDF15. Together, these findings suggest GDF15 can be used to study the energetic mechanisms for the biological embedding of stress across the lifespan.

## 1. Introduction

People experiencing higher levels of psychosocial stress exhibit accelerated biological aging and elevated chronic disease risk(1). However, much remains unknown about molecular pathways linking stress to aging. Stress requires energy, and alterations in energy metabolism contribute to the biological embedding of stress(2). Emerging evidence suggests that psychological stress alters mitochondrial biology and bioenergetics(3–7), which are central regulators of both aging(8–13) and disease processes(14, 15). Together, this evidence supports a pathway in which mitochondria serve as a key link between stress, aging, and disease. While preclinical models have begun to elucidate how psychological stress affects mitochondrial biology leading to accelerating biological aging and increasing vulnerability to chronic diseases(16, 17), its translation to humans remains underexplored.

To conduct analysis of the association between psychological stress and mitochondrial biology in humans, we focused on a circulating biomarker of energetic stress, the protein Growth Differentiation Factor 15 (GDF15)(12). GDF15 is a cytokine produced by various tissues and secreted in the bloodstream. It is hypothesized to signal energetic stress to the brain, influencing behavior and energy expenditure(18–22). Circulating GDF15 levels are elevated in mitochondrial energy transformation disorders (i.e., mitochondrial diseases)(23–25) and increase with nutritional and physiological stressors such as exhaustive exercise(26, 27), pregnancy(19, 28), multi-day starvation(29), and infectious challenges(30). These changes in circulating levels of GDF15 are meaningful for human health, as indicated by the robust elevation of GDF15 with aging(31, 32) and its associations with the risk for many aging-related chronic diseases and mortality(31, 33–40). However, whether GDF15 is associated with psychological stress in humans remains unclear.

To address this knowledge gap, we conducted a series of epidemiological and experimental studies. First, using data from the Framingham Heart Study and UK Biobank, we tested whether plasma GDF15 levels were elevated in individuals exposed to chronic psychological stressors, including mood disorder symptoms, low educational attainment, low family income, and high job strain. Second, using intensive sampling for two individuals over 53 and 60 consecutive days, we investigated whether saliva GDF15 exhibits an awakening response, a dynamic and actively regulated pattern characteristic of other stress hormones(41, 42). Finally, in two laboratory experiments, we tested whether acute psychosocial stress induced changes in GDF15 levels in plasma and saliva. Together, our findings establish a foundation for studying GDF15 as a novel biomarker at the interface of psychosocial stress, mitochondrial biology, and aging.

## 2. Results

### Blood GDF15 and mental health

Psychological stress is a well-established contributor to the onset and expression of mental disorders(43, 44). As an initial test to evaluate whether GDF15 is linked to psychosocial stress, we examined its association with mental health symptoms using data from the *Framingham Heart Study, Offspring cohort* (*FSH,* n=3,460) and with mental health disorder diagnosis using the proteomics atlas(38) of *UK Biobank* (*UKB*, n=53,026) (Figure 1A). Given the evidence for sex differences in mental health symptoms(45), we analyzed associations separately for women and men(46).

**Figure 1.**
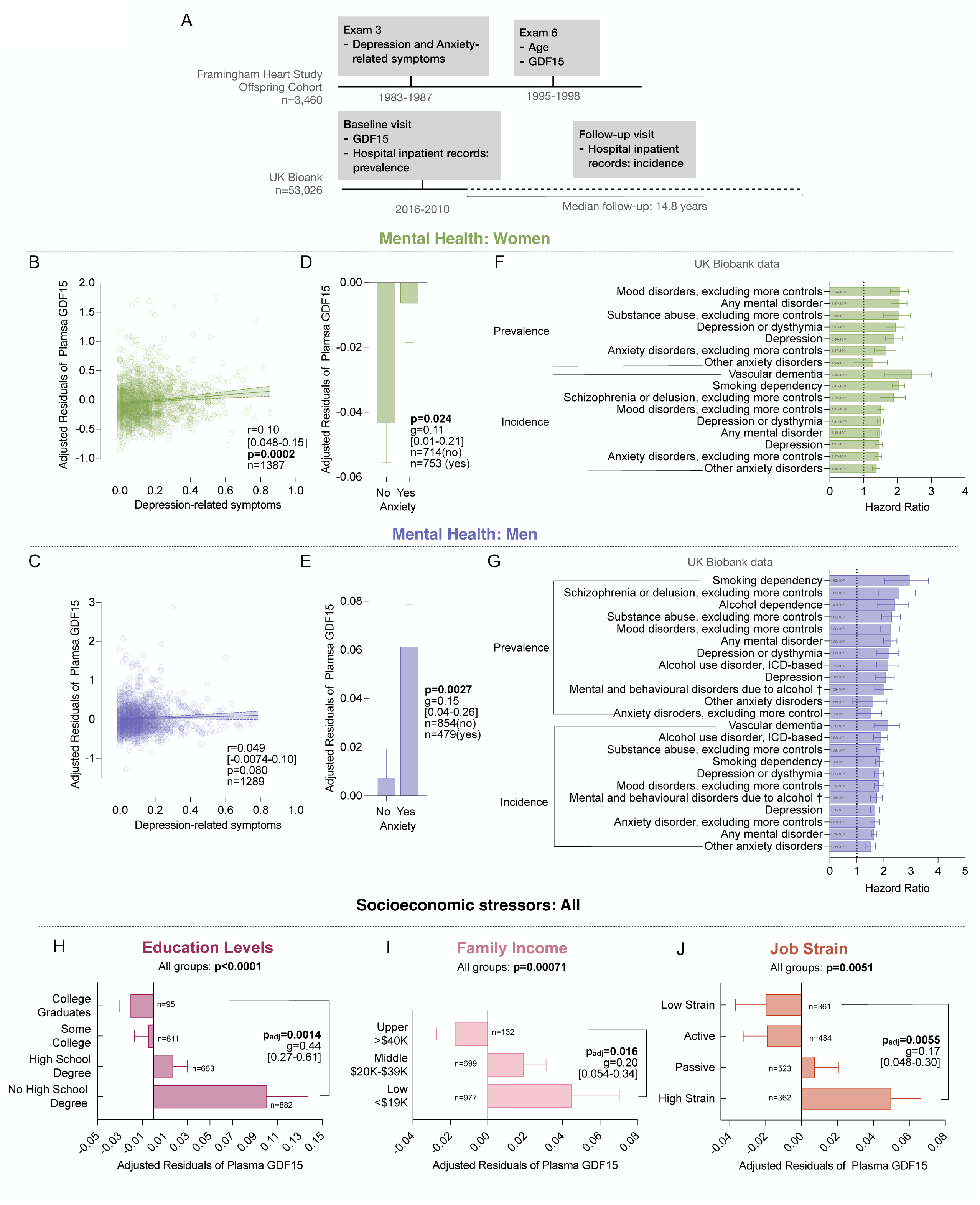
Associations of plasma GDF15 and epigenetic clocks with depression, anxiety, and mental illness prevalence and incidents. (**A**) FHS and UKB designs. (**B,C**) Scatterplot of the associations between age-adjusted GDF15 levels and depression-related symptoms (FHS) for (**B**) women and (**C**) men, see Supplemental Figure S1A for results from combined data for women and men analysis). (**D,E**) Average GDF15 levels by anxiety-related symptoms for (**D**) women and (**E**) men in FHS (Supplemental Figure S1B for women and men combined analysis). (**F,G**) Associations between GDF15 levels and mental disorder incidence and prevalence (UKB) for (**F**) women and (**G**) men. UKB GDF15 data measured using Olink proteomics and modeled as a continuous variable using NPX values where each 1-unit increase reflects a doubling in protein concentration; hazard ratios represent the change in risk per doubling of GDF15. (**H-J**) Average GDF15 by (**H**) education levels (n=2191), (**I**) family income levels (n=1809), (**J**) job strain (n=1730) defined as 4 categories: Low Strain (high in decision latitude and low on psychological job demand); Active (high on both decision latitude and psychological job demand); Passive (low on both decisions attitude and psychological job demand); High Strain (Low on decision latitude and high on psychological job demand). P-values from (B,C) Spearman’s rho [95% confidence interval], (D,E) Mann-Whitney rank t-test, (H-J) Krusakal-Wallis test and Dunn’s multiple comparison test adjusted for multiple testing. (D,E,H-J) Data shown as mean ± SEM, effect sizes estimated using Hedges’ g [95% confidence interval]. (F,G) Logistic regression and Cox proportional hazards models were used to assess GDF15’s associations with prevalent disease and incident diseases, respectively. In UKB, GDF15 was modeled as a continuous predictor using Normalized Protein eXpression (NPX) values (log_₂_-transformed, normalized protein abundance). A 1-unit increase in NPX reflects a doubling of GDF15, therefore hazard ratios represent risk per doubling of its concentration. All regressions we’re performed adjusting for baseline age, sex, ethnicity, Townsend deprivation index, BMI, smoking status, fasting time, season of blood collection, and blood age (date of blood collection to date of protein examination) as covariates. Association with depression and anxiety-related diagnosis, as well as other mental health-related diagnosis that passed Bonferroni correction (p=1×10-9) were presented in graphs. Data shown as hazard ratio ± 95% confidence interval, p-value shown annotated in each bar. † excluding acute intoxication.

In *FHS*, we tested cross-sectional associations between depressive and anxiety symptoms reported by participants during the third examination cycle and plasma GDF15 levels measured in pg/mL from blood draws at the sixth examination cycle. Depressive symptoms were positively associated with age-corrected GDF15 levels observed in women (Figure 1B, Spearman’s r=0.10, 95% Confidence Interval [0.048-0.15], p=0.0002). In men, we observed a similar direction of association, but the magnitude was smaller and not statistically different from zero at the α=0.05 level (Figure 1C, r=0.049, 95% CI [-0.0074-0.10], p=0.080). Similarly, participants with anxiety symptoms exhibited higher age-corrected GDF15 levels than those without symptoms, in both women (Figure 1D, Hedges’ g=0.11, 95% CI [0.01-0.21], p=0.024) and men (Figure 1E, g=0.16, 95% CI [-0.0074-0.10], p=0.0027).

In the *UKB*, we examined differences in GDF15 levels between individuals with and without prevalent mental disorders. We also explored whether baseline GDF15 levels were prospectively associated with incident of mental disorders over 15 years of follow-up. GDF15 was modeled as a continuous predictor using Normalized Protein eXpression (NPX) values, which are log_₂_-transformed and normalized measures of protein abundance. A 1-unit increase in NPX corresponds to a doubling of GDF15 levels; thus, hazard ratios represent the risk per doubling of GDF15 levels. At baseline, higher levels of blood GDF15 were associated with greater prevalence of both depression (for women Hazard Ratio=1.91, 95% CI [1.68-2.18], p=3.9×10^-22^; for men HR=2.06, 95% CI [1.81-2.60], p=6.1×10^-17^) and anxiety disorder (for women HR=1.67, 95% CI [1.38-2.02], p=1.4×10^-7^; for men HR=1.53, 95% CI [1.14-2.05], p=4.4×10^-3^). Over follow-up, those with higher GDF15 levels were more likely to have an incident diagnosis of both depression (for women HR=1.46, 95% CI [1.37-1.56], p=1.3×10^-30^; for men HR=1.68, 95% CI [1.53-1.85], p=2.8×10^-27^) and anxiety disorder (for women HR=1.44, 95% CI [1.34-1.55], p=1.6×10^-23^; for men HR=1.66, 95% CI [1.49-1.84], p=4.8×10^-21^). Results are shown in Figures 1F-G. Together, our findings showing the association between elevated GDF15 and mental health symptoms confirm previous studies linking mood disorders with elevated GDF15(35, 47–49) and suggest that elevated GDF15 could contribute to the development of mental illness.

### Blood GDF15 and socioeconomic stressors

To test whether GDF15 reflects the broader, cumulative burden of psychosocial stress beyond clinical symptoms, we analyzed associations with educational attainment, family income, and job strain. We report effect-sizes in terms of GDF15 standard deviation units in the *FHS*.

In *FHS*, participants who completed higher levels of education had lower GDF15 levels compared to those with lower levels of education (Figure 1H, college graduates vs no high school degree: g=0.44, 95% CI [0.27-0.61], p_adj_=0.0014). Findings were similar in *UKB* where more years of education was associated with lower levels of GDF15 (unstandardized β=-0.28, SE=0.068, 95% CI [−0.41 to −0.15], p<0.001).

*FHS* participants reporting higher family income had lower GDF15 levels compared to those reporting lower incomes (Figure 1I, upper vs low income class: g=0.20, 95% CI [0.054-0.34], p_adj_=0.031).

*FHS* participants reporting high job strain had higher GDF15 levels compared to those reporting low job strain (Figure 1J, high vs low job strain: g=0.17, 95% CI [0.048-0.30], p_adj_=0.031). Together these findings suggests that psychosocial stress is associated with higher levels of GDF15.

### Benchmarking stress-GDF15 associations against epigenetic clocks

Psychosocial stressors such as mental health symptoms and socioeconomic factors are hypothesized to accelerate biological aging(50). Consistent with this hypothesis, people with histories of mental health problems and socioeconomic disadvantages show signs of accelerated biological aging as reflected in a family of DNA methylation (DNAm) algorithms known as epigenetic clocks(51–53). To provide context for the effect-sizes we observed for GDF15 and psychosocial stressors, we repeated our *FHS* analysis with epigenetic clock measurements made from blood DNA collected at the 8^th^ examination cycle (Figure 2A).

**Figure 2.**
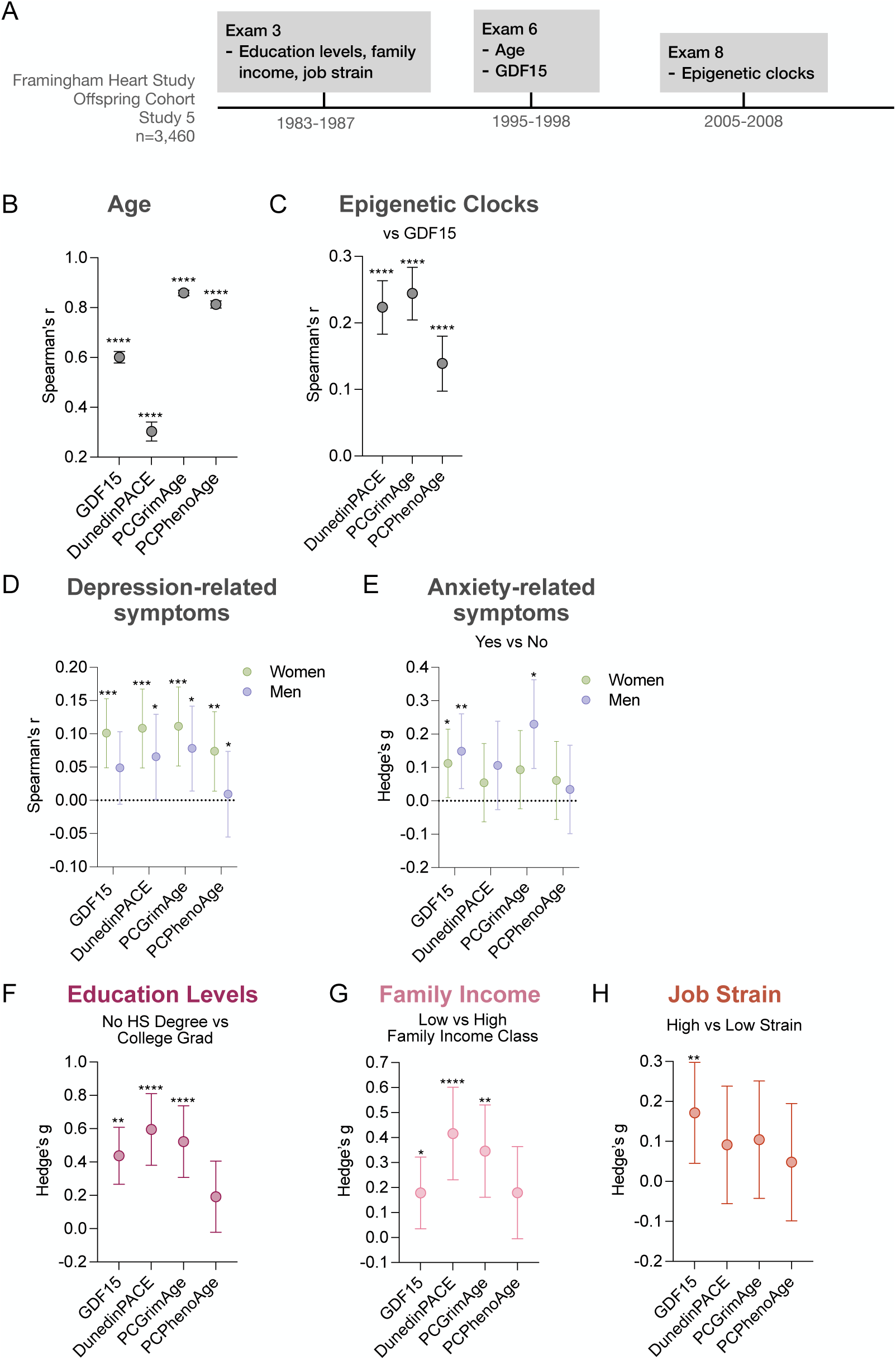
Associations of plasma GDF15 and epigenetic clocks with socioeconomic and aging factors. (**A**) FHS design. (**B**) Effect sizes for the associations between age, GDF15 (n=2,900), and epigenetic clocks (n=2,160), estimated using Spearman’s rank correlation. See Supplemental Figure S1C for individual scatterplots. (**C**) Effect sizes for the associations between age-corrected plasma GDF15 levels and each epigenetic clock (n=2,160), estimated using Spearman’s rank correlation. See Supplemental Figure S1D for individual scatterplots. (D,E) Effect sizes for the associations between plasma GDF15 levels, epigenetic clocks and (D) depression-related symptoms and (E) anxiety-related symptoms; shown for women and men separately (see Supplemental Figure S2 for analysis of each epigenetic clock, presented separately for women and men).(F-H) Effect sizes (Hedge’s g and 95% CI from normal approximation) comparing average plasma GDF15 levels and epigenetic clocks in participants: (F) with no high school degree vs. college degrees; (G) in low vs high family income class; (H) with low vs high job strain. See supplemental figure S3 for average epigenetic clock value by education, family income, and job strain for DunedinPACE, GrimAge, PhenoAge. (B-D) Data shown as mean ± SEM. P-values from Krusakal-Wallis test and Dunn’s multiple comparison test adjusted for multiple testing. Effect sizes from (E,F) Spearman’s rho and (G-I) Hedges’ g. (E-I) Data shown as ± 95% confidence interval calculated from (E,F) Fisher’s Z-transformation and (G-I) normal approximation. Analysis performed on (B,C,D,F,G,H,I) age- and sex-adjusted and on (B) raw GDF15 and epigenetic clocks values. *p<0.05, **p<0.01, ***p<0.001, ****p<0.0001.

We analyzed the three epigenetic clocks(54, 55) with the strongest validation track record in studies of healthspan and lifespan, PhenoAge(56), GrimAge(57), and DunedinPACE(58–60). GDF15 and the epigenetic clocks were correlated with age (r=0.3-0.8) and, after correction for age differences, with each other (r=0.14-0.24) (Figures 2B-C). Effect sizes for GDF15 associations with mental health and socioeconomic outcomes were similar to those for epigenetic clocks, with the exception that GDF15 associations with job strain, were somewhat stronger as compared to the clocks (Figure 2D-G). GDF15 was uniquely associated with job strain (Figure 2H), suggesting that this biological aging marker might be closely linked to this stressor than the epigenetic clocks. Taken together, our results corroborate previous research(61) showing that psychosocial stressors are linked to accelerated aging as measured by epigenetic clocks. Furthermore, these results suggest that GDF15 may be similarly effective or better in capturing these stress to accelerated aging associations.

### Dynamic Variation of Saliva GDF15

Our epidemiological findings suggest that people exposed to chronic stress have elevated levels of GDF15. We previously demonstrated that GDF15 can be measured in saliva and is elevated in individuals with mitochondrial DNA defects(62). We examine whether GDF15 is dynamically regulated in the context of a physiological stressor known to affect other biomarkers of psychological stress: the awakening response. Waking up is a metabolically demanding transition(63) associated with cortisol awakening response (CAR), which reflects hypothalamic-pituitary-adrenal (HPA) axis activity and predicts stress dysregulation conditions such as depression(64). By testing whether saliva GDF15 exhibits an awakening response, we aim to investigate whether saliva GDF15 exhibits dynamic properties and responds to acute physiological demands of daily life.

To explore potential diurnal variations, we first examined serum GDF15 levels across waking and sleep hours and found no systematic pattern (see SI Text 1.1 and SI Text Figure 1 for additional details). We then assessed saliva GDF15 levels from an *Intensive repeated-measures n=2 study*, where two male participants were sampled over 53-60 consecutive days. Using a home-based collection designed to detect awakening responses, participants provided four samples per day: at awakening, +30 min, +45 min, and bedtime (Figure 3A). Saliva GDF15 exhibited a robust negative awakening response, declining on average by 61% and 89% at 45 min in Participant A and B, respectively (Figures 3B-C). These intensively sampled measurements provided preliminary evidence on important within-individual variation (+140% to −98% at 45min depending on the days) in saliva GDF15 dynamics (Figures 3D-E). See SI Text 1.2 and SI text Figure 2 for additional details.

**Figure 3.**
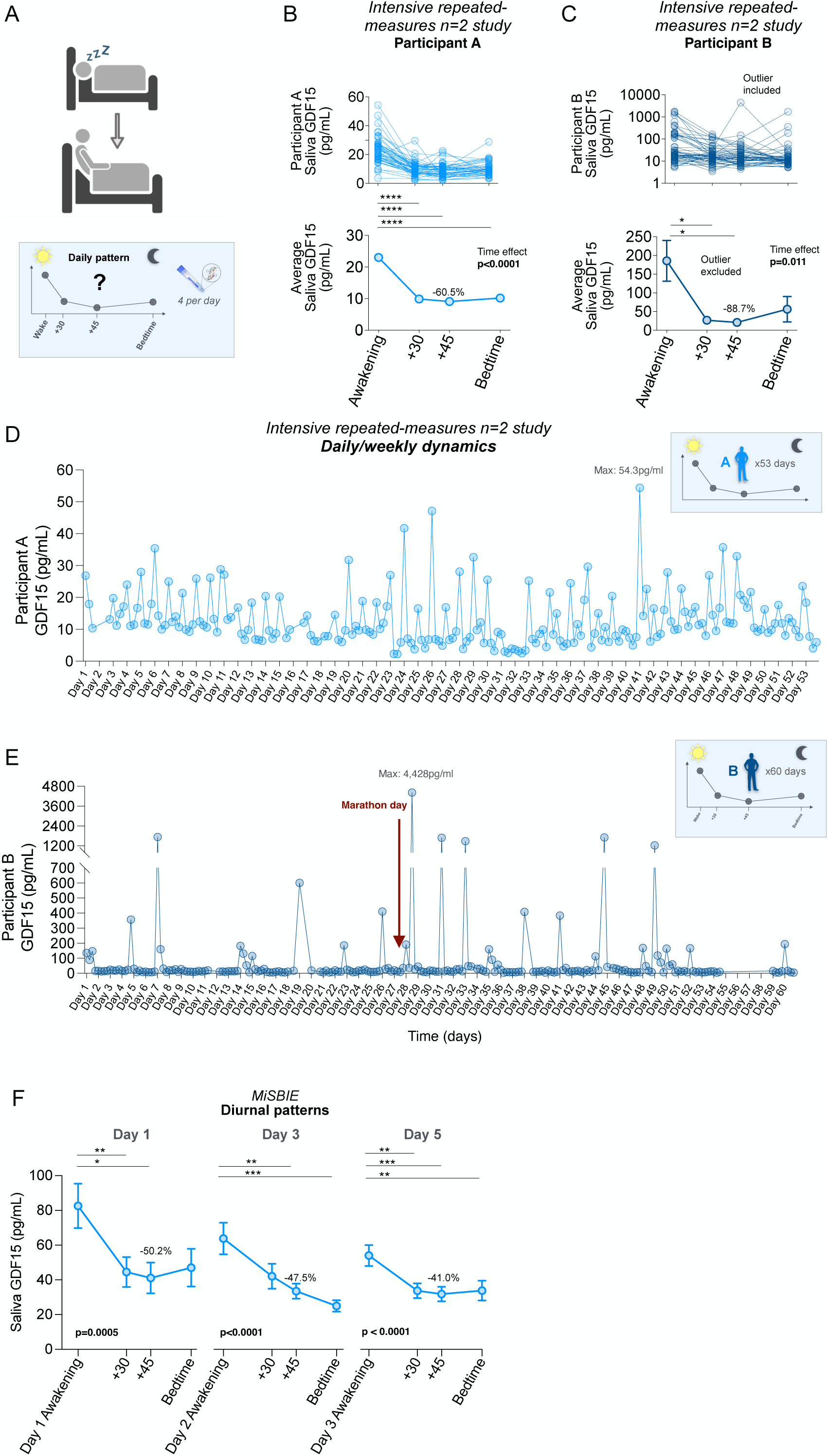
Saliva GDF15 exhibits a robust negative awakening response and large within-person variation over days to weeks. (**A**) At-home awakening response study design measuring saliva GDF15. Saliva was sampled at awakening, 30 and 45 min after awakening, and at bedtime. (**B,C**) Saliva GDF15 diurnal variation from intensive repeated-measures n=2 studyParticipant A (**B**) and B (**C**). Top, individual days; bottom, average trajectory. One outlier timepoint, Day 28, 45 min, was removed from analyses. See figure S4A for median of diurnal pattern for Participant A and B. (**D,E**) Day-to-day variation of saliva GDF15 levels from Participant A (**D**, n=199 observations, repeated measures for 53 days) and Participant B (**E**, n=221 observations, repeated measures for 60 days).(**F**) Saliva GDF15 diurnal variation from MiSBIE healthy adults on three non-consecutive days four timepoints each day (n=710 observations, 68% female), see Supplemental figure S4B for individual trajectory. Data shown as (B,F,G) mean ± SEM and (B) median ± 95% Cl. Effect sizes and P-values from (B,C,F) Mixed-effect model and Tukey multiple comparison test corrected for multiple testing. Saliva data below minimum detection value were included in graphing but excluded in statistical analysis. *p<0.05, **p<0.01, ***p<0.001, ****p<0.0001.

We validated this robust negative awakening response in the *Stress, Brain Imaging, and Epigenetic Study (MiSBIE)*(*65*), where 70 healthy individuals (mean age 37.1 years, 68% female) collected saliva using the same protocol for 3 non-consecutive days at home. Similar to what was found in the *Intensive repeated-measures n=2 study*, GDF15 was highest immediately at the time of waking up (i.e., awakening) before sharply declining by an average of −40% at 30 min, and −46% at 45 min, reflecting a robust negative awakening response (Figure 3F). At bedtime, saliva GDF15 was, on average, 47% lower than the awakening value. We observed important between-individual variation in saliva GDF15, with some participants’ levels increasing by up to 463% and others declining by up to 97% at 45min.

In *Intensive repeated-measures n=2 study* and *MiSBIE*, we assessed salivary cortisol levels and found no association between the magnitude of GDF15 and the cortisol awakening response in either study (Figures S4C-D).

Together, these findings demonstrate that saliva GDF15 exhibits a robust awakening response, like other neuroendocrine factors(66). In stress research, the CAR has been a valuable tool in studying HPA axis dysregulation in relation to chronic stress or mood disorders(67, 68). The important within- and between-individual variation in the magnitude of the GDF15 awakening response could be further explored in relation to chronic stress. These data lay the groundwork for future studies to investigate saliva GDF15 as a potential biomarker of mitochondrial energetic stress in relation to daily stress.

### Psychobiological regulation of plasma and saliva GDF15

Having established that elevated GDF15 levels are associated with chronic psychological stressors and that GDF15 exhibits a robust awakening response, we next sought to determine whether GDF15 is responsive to acute psychological stress. To conduct this analysis, we analyzed data collected within the *MISBIE* study using an experimental paradigm that is well established in stress biology: a 5 min social-evaluative speech task(69).

Participants (70 healthy individuals, mean age 37.1 years, 68% female) were asked to defend themselves against accusation of an alleged transgression. This social-evaluative stress task included 2 minutes of preparation and three minutes for speech delivery in front of a white coat-wearing “evaluator”, a mirror, and a camera. Plasma and saliva were concurrently collected at eight timepoints: 5 min before and 5, 10, 20, 30, 60, 90, 120 min after the initiation of the stressor (Figure 4A).

**Figure 4.**
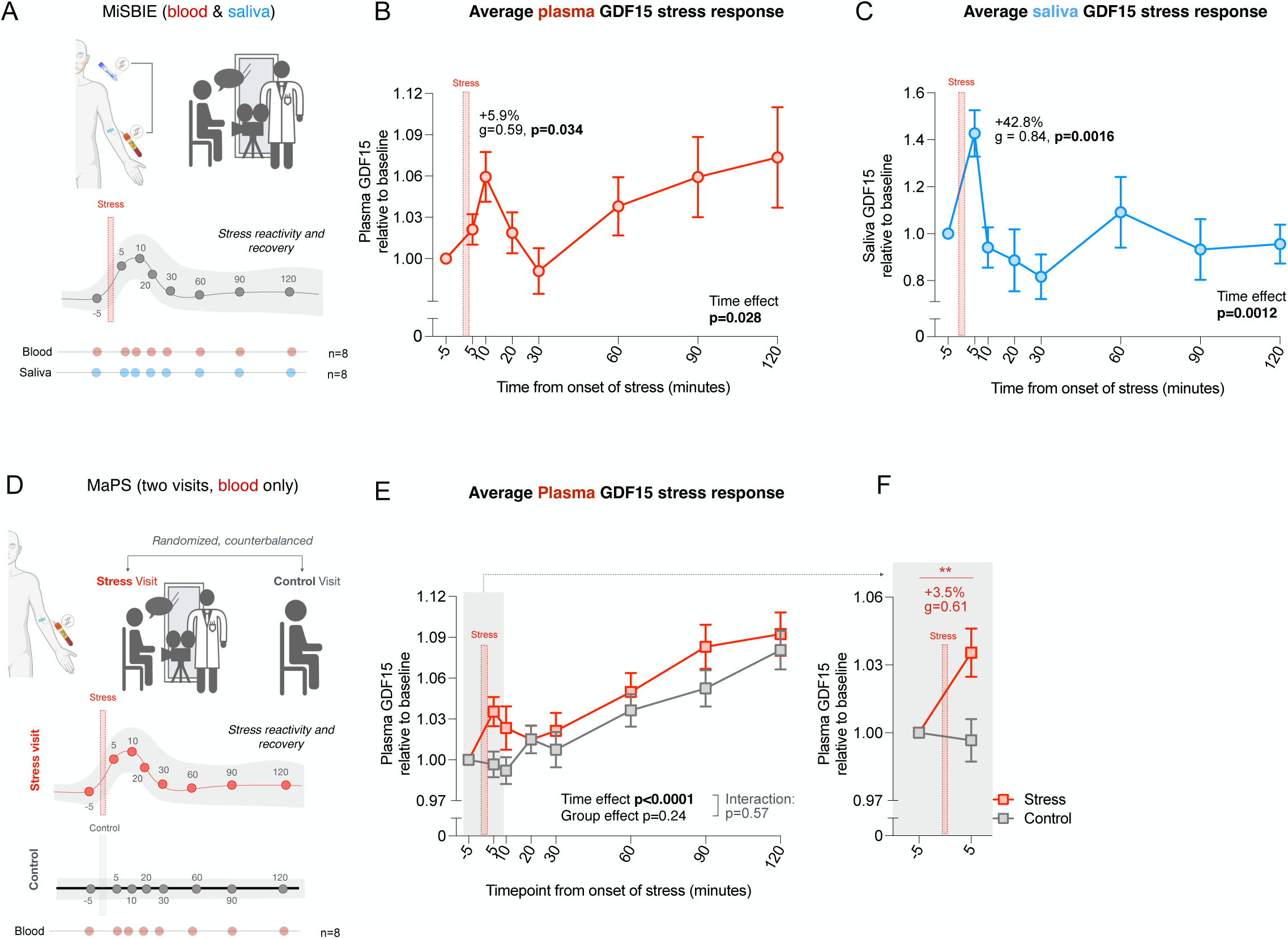
Social-evaluative stress acutely increases plasma and saliva GDF15. (**A**) MiSBIE study design measuring psychological response in GDF15 after social-evaluative stress elicited by a validated social-evaluative speech task.(**B,C**) Plasma (**B**) and Saliva (**C**) GDF15 levels before and after psychological stress shown as percent change from baseline (−5 min). For plasma n=65, 68% female, 512 observations, for saliva GDF15 n=57, 65% female, 442 observations. For individual trajectories and absolute change see Supplemental Figure S5A-D.(**D**) MaPS study design including two visits randomized to either social-evaluative speech task (same as in MiSBIE) or a control session without stress exposure. Plasma GDF15 was measured at 8 timepoints before and after stress, following the same schedule as MiSBIE. (**E**) Plasma GDF 15 levels for control and stress visit shown as percent change from baseline (−5min). n=70, 48% female, 916 observations. For individual trajectories and absolute change see Supplemental Figure S5E-F. (**F**) Plasma GDF15 reactivity focused on −5 vs 5min timepoints for control and stress visit shown as percent change from baseline, p-values from Wilcoxin paired t-test for stress group, effect size from Hedges’ g comparing −5 to 5min for stress group. n=58 pairs, 48% female, 117 observations. (B,C,D,F) Data shown as mean ± SEM. P-values from (B) Wilcoxon paired t-test, (C,D) mixed-effect model followed by Tukey multiple comparison test, corrected for multiple testing, (E) two-way ANOVA followed by Tukey multiple comparison test, corrected for multiple testing. *p<0.05, **p<0.01, ***p<0.001, ****p<0.0001.

At baseline, 5 min prior to the initiation of the social-evaluative challenge, participants had plasma GDF15 concentrations of M=385.27pg/mL (SD=157.77). In response to the challenge, GDF15 levels increased by 5.9% ([95%CI 2.3%-9.0%], p=0.034) across the first 10 min after the stressor before declining to baseline levels by the 30 min follow-up. GDF15 concentrations in saliva at baseline were M=23.39pg/mL (SD=24.27). Saliva GDF15 also responded to the stressor, peaking 5 min after initiation of the challenge, and declining to baseline levels by the 10 min follow-up. Strikingly, the saliva response was much larger; GDF15 levels rose by 43% ([95%CI −2.3%-63%], p=0.0016), a nearly seven-fold larger response as compared with plasma GDF15. Plasma- and saliva-GDF15 trajectories are shown in Figure 4B-C. This represents, to our knowledge, the first evidence that mental stress alters GDF15 concentrations in accessible biofluids within minutes in human.

To validate the speech-task-induced changes in plasma GDF15, we analyzed samples from *MaPS* (*Mitochondria and Psychological Stress Study*) where 68 healthy individuals (mean age 30.6 years, 51.2% female) completed two laboratory visits in randomized order, involving either the same speech task as in *MiSBIE* or a control condition (no stressor beyond the blood draw and laboratory setting) (Figure 4D). Plasma GDF15 was measured at the same 8 timepoints as in *MiSBIE*. *MaPS* validated the direction and magnitude of results from *MiSBIE* where mental stress increased plasma GDF15 by 3.5% ([95%CI 1.4%-5.7%], p=0.0085) at the 5 min timepoint (Figures 4E-F). See SI Text 1.3 and SI Text Figure 3 for additional details.

In *MiSBIE*, plasma and saliva GDF15 levels both responded to the social-evaluative stress task; but the responses within each person were not correlated, potentially pointing to an independent (psycho)biological regulation of GDF15 across biofluids (Figure S6A). In *MiSBIE and MaPS*, we also assessed levels of cortisol, norepinephrine, and epinephrine during the social-evaluative stress task. No correlation was found between the magnitude of the plasma or saliva GDF15 response and the responses of these stress hormones (see Figures S6B-G for *MiSBIE* results, see SI Text 1.3 for *MaPS* results).

To compare GDF15 to another energetic stress marker, lactate (produced by increasing the intracellular NADH/NAD^+^ ratio(70)), we explored lactate stress reactivity in a subset of the *MiSBIE* participants. Similar to what was previously observed in mice(71), evoked mental stress elevated circulating lactate in humans (see SI Text 1.4 and SI Text Figure 4 for additional details). In relating GDF15 and lactate, we found a modest positive correlation between the saliva GDF15 response and the acute rise in blood lactate (r=0.32, p=0.045; Figure S6H), consistent with a potential link between systemic reductive stress and saliva GDF15 reactivity.

## 3. Discussion

We conducted a multi-study, multi-method investigation of the circulating cytokine GDF15 as a peripheral biomarker of psychosocial stress. There were four main findings. *First*, people exposed to chronic stressors had higher levels of GDF15. In two prospective cohort studies, we observed elevated plasma GDF15 levels in individuals reporting symptoms of depression and anxiety, lower socioeconomic status as reflected in their income and educational level, and higher levels of job strain. *Second*, people with elevated plasma GDF15 levels show signs of accelerated biological aging. In a prospective cohort study, participants with higher GDF15 levels later exhibited faster pace of aging and older biological age as measured by epigenetic clocks. *Third*, saliva GDF15 is dynamically regulated by acute physiological stress. In two independent studies, we observed peak levels of saliva GDF15 upon waking up, with declines to a stable minimum within 30-45 minutes that was maintained through bedtime. *Fourth*, saliva and blood GDF15 are dynamically regulated by acute psychological stress. In two social-evaluative stress experiments, participants’ GDF15 levels increased within minutes of stress induction. We observed substantial individual differences in the magnitude of the GDF15 awakening response, as well as in the recovery time and trajectory of the GDF15 response to acute psychological stress. Together, these findings: 1) link GDF15 to chronic psychosocial stress and accelerated biological aging; 2) establish plasma and saliva GDF15 as dynamic markers responsive to short-term physiological and psychological stress; and 3) highlight its potential as a biomarker of psychosocial stress.

Our findings of higher GDF15 levels in people with depression and anxiety symptoms expand on previous findings showing elevated GDF15 levels in psychiatric diseases(35, 47–49). The prospective association found between elevated GDF15 levels and greater risk of anxiety and depression after a 15-years-follow-up in the *UKB* study supports the emerging notion that psychiatric disorders may arise, in part, from stress-induced energetic dysregulations(72, 73).

Similar to immune-derived cytokines that trigger sickness behavior by neural signaling(74), GDF15 may similarly act as a brain-acting signal(20), inducing energy conservation behaviors(22) to meet stress-related energy demands and promote resilience. However, prolonged activation of this pathway could lead to psychiatric symptoms such as depression and anxiety(12, 75). Findings from animal models also support GDF15’s role in the regulation of anxiety-like behavior: GDF15 knockout mice exhibit reduced anxiety(76), while elevated GDF15 induced by muscle mitochondrial dysfunction(77), exogenous administration of GDF15(78), or stressors such as adrenaline injection and cage change(79) leads to increased anxiety-like behavior, likely mediated via GDF15-GFRAL signaling. Collectively, these findings suggest a biological pathway in which psychological stress elevates GDF15, potentially increasing vulnerability to psychiatric symptoms through GDF15-GFRAL signaling in the central nervous system. Further longitudinal studies are needed to study the bioenergetic contribution of mental health disorders(80) in greater depth.

A key feature of a robust stress biomarker is its ability to reflect the cumulative burden of chronic psychosocial stress, such as low socioeconomic status. Our findings of higher GDF15 in people exposed to socioeconomic disadvantages expend prior observations linking elevated GDF15 to lower education attainment(81) and poverty status(82). We replicated these associations in two large cohorts and further demonstrated similar patterning of GDF15 with family income and job strain. Importantly, when benchmarked against epigenetic clocks, validated biomarkers of biological aging, GDF15’s associations with psychosocial stressors showed effect sizes of comparable or better magnitude. These findings underscore the sensitivity of GDF15 to capture the biological impact of socioeconomic disadvantage and psychosocial stress.

Our findings showing that plasma GDF15 levels are positively associated with epigenetic clocks suggest that individuals experiencing cellular energetic stress (indexed by higher GDF15) exhibit accelerated biological aging. This aligns with previous findings showing that greater GDF15 is associated with accelerated aging indexed by epigenetic clocks(83) and the Pace of Aging composite score(84). Framed within emerging work positioning mitochondria as a driver of both diseases(17) and aging(8–11), GDF15’s association with mental health, socioeconomic stressors, and epigenetic clocks suggest that *Stress* _→_ *Aging*_→_ *Disease pathway* may include cellular energetics as a key mediator. Within that framework, prolonged exposure to socioeconomic disadvantage might increase the energetic burden of the organism, accelerate physiological aging, and increase vulnerability to diseases(85, 86). Future studies should test whether stress-induced elevations in GDF15 are associated with subsequent adverse aging trajectories.

While accumulating evidence show a connection between elevated GDF15 and health trajectories(37), little is known about the dynamic inducibility of GDF15 by acute physiological and psychological stress. The robust decline in saliva GDF15 we observed within 30 to 45 min after waking illustrates its capacity for rapid fluctuations after physiological stress similar to other stress biomarkers(87, 88). Our findings also demonstrate that GDF15 responds to a brief and relatively mild 5 min social-evaluative stressor. Across two independent studies, plasma GDF15 increased modestly (+3–6%), while saliva GDF15 exhibited a much larger response (+42–71%). Notably, both plasma and saliva levels rose within 5–10 min and declined within 30 min on average, demonstrating a rapid and dynamic response to acute stress. Extensive research on cortisol has shown that variations in the pattern of its awakening response and acute responses to psychosocial stress are associated with both stress exposure (e.g., childhood trauma(89, 90)) and stress-related conditions (e.g., depression(64, 91)), contextualizing the role of HPA axis in the biological embedding of stress. Similarly, our finding that GDF15 exhibits a robust awakening response and acute psychological stress reactivity, along with substantial individual differences in response magnitude, recovery time, and trajectory, opens new avenues for investigating the role of energetic stress signaling in connection to psychological stress and stress-related conditions.

Preclinical models suggest a complex, potentially bidirectional interaction between GDF15 and stress hormones. Exogenous administration of adrenaline administration increases circulating GDF15(79), while GFRAL+ neuron activation elevates circulating catecholamines(22). Exogenous administration of GDF15 increase elevate circulating corticosterone(79, 92), while endogenous GDF15 is required for full HPA axis activation in response to toxins that do not induce a substantial cytokine response(92). In humans, cortisol deficiency has also been associated with elevated GDF15 levels(93). The apparent lack of association between GDF15, cortisol, and catecholamine responses to acute physiological and psychological stress in our human studies requires further investigation. Functional interactions between the HPA and the ISR-GDF15 axis might involve complex dynamics, such as compensatory mechanisms, that need to be further studied in humans.

Several limitations and unresolved questions should be noted. First, the *FHS* and *UKB* cohorts are not population representative studies. They over-represent white and socioeconomically better-off segments of the population. Future studies should aim to replicate findings in other populations and in more representative cohorts. Second, a common GDF15 polymorphism (H202D, present in 15-30% of the population(94)) has been reported to reduce GDF15 detection by the R&D ELISA kits(95). The R&D ELISA kits were used to assess GDF15 concentrations in plasma and saliva samples from several of our studies (*MiSBIE*, *MaPS*, *Intensive repeated-measures n=2 study*, and *overnight meal alignment study*). While this may affect between-group comparisons, within-person analyses of awakening responses and stress reactivity would not be affected. Future study should account for this assay limitation when designing protocols. Finally, the lack of correlation between plasma and saliva GDF15, along with the significantly larger saliva response (+42.8% vs. +5.9% in plasma) require further investigation by future studies. GDF15 is expressed by salivary glands(62), suggesting an independent regulation from blood levels and its secretion into saliva might play a role in wound healing(96).

In summary, our findings reveal that GDF15 is associated with chronic and acute psychosocial stress, establishing GDF15 as a dynamic biomarker at the intersection of psychosocial stress, energetic stress, and biological aging. Future research should investigate GDF15 secretion and signaling mechanisms, its role in stress-related diseases and aging biology, and its potential as a tool monitoring how both negative and positive experiences become biologically embedded across diverse populations. Ultimately, our research positions GDF15 not only as a promising biomarker for psychosocial stress but also as a potential target for interventions to mitigate the adverse effects of chronic stress on mental health and biological aging, thereby paving the way for future studies that could transform our understanding of stress-related diseases.

## 4. Methods

### 4.1. Framingham Heart Study

#### 4.1.1 Participants and procedures

The *Framingham Heart Study (FHS)*, launched in 1948 in Framingham, Massachusetts, has tracked cardiovascular disease progression across three generations(97). Our analysis focused on data from the Offspring Cohort—the study’s second generation—recruited from 1971. We analyzed data from psychosocial questionnaire package collected during the cohort’s third examination cycle (1983–1987), plasma GDF15 measured from blood samples collected during the cohort’s sixth examination cycle (1995–1998), DNA methylation analysis from blood samples collected during the cohort’s eighth examination sample (2005–2008).

#### 4.1.2 Psychosocial questionnaire package

A 300-item questionnaire was developed to assess five key aspects of psychosocial stress: socio-demographics, life events, behavior types, situational stress, and somatic strain(98). In this study, we analyzed questions related to education attainment, family income, job strain, depression-related symptoms, and anxiety-related symptoms. Education attainment was assessed categorically: 1: No High School Degree; 2: High School Degree; 3: Some College; 4: College Graduate. Family income was assessed categorically: 1: no income; 2: less than $5,000; 3: $5,000 to $9,000; 4: $10,000 to $14,000; 5: $15,000 to $19,000; 6: $20,000 to $24,000; 7: $25,000 to $29,000; 8: $30,000 to $34,000; 9: $35,000 to $39,000; 10: $40,000 to $44,000; 11: $45,000 to $49,000; 12: more than $50,000. For this study, we grouped family income responses into three categories based on 1980s income cut-off: Low <$19K; Middle $20K −$39K; High >$40K. Job strain questionnaire was adapted from Robert Karasek’s Demand-Control Model of Job Stress(99). Questions were grouped as Decision Latitude Scale and Psychological Job Demand and required Likert-type (very well, fairly well, somewhat, not at all) responses. Based on their responses, participants fell into 4 categories: 1: High Strain, based on simultaneously high job demands (above the median score) and low job decision latitude (at or below the median score); 2: Low Strain, based on simultaneously low job demands and high job decision latitude; 3: Passive, based on simultaneously low job demands and low decision latitude; or 4: Active, based on simultaneously high job demands and high job decision latitude. Depression-related symptoms were measured as a continuous scale from 0 to 0.85, with 0.85 being the most depressed. Anxiety-related symptoms were assessed using five questions requiring dichotomous (yes or no) responses. Participants were grouped as ’no’ if they answered ’no’ to all five questions and as ’yes’ if they answered ’yes’ to at least one.

#### 4.1.3 Plasma GDF15

At the sixth examination cycle, circulating levels of GDF15 was assessed alongside a panel of other biomarkers as detailed previously(100). Briefly, blood samples were collected following an overnight fast, then promptly centrifuged and stored at −80°C until analysis. GDF15 levels were determined using a precommercial immunoassay on a Cobas e 411 analyzer (Roche Diagnostics, Switzerland).

#### 4.1.4 Epigenetic clocks

DNA methylation data was analyzed from blood samples collected in 2,471 Framingham Study participants (54% female, mean age 66, SD = 9) at the eighth examination cycle. Whole-genome DNA methylation data were accessed from the NIH database of Genotypes and Phenotypes (dbGaP; accession number phs000724.v9.p13). Informed consent was not required, as deidentified data from the FHS were used. Detailed processing and analysis can be found in previous publications(61, 101). In brief, methylation was measured using Illumina 450K arrays (2005–2008) and processed by the Columbia Aging Center, yielding quality-controlled data for 2,296 participants. Epigenetic clocks (DunedinPACE(102), PC version of GrimAge(57), PC version of PhenoAge(56)) were calculated with established R packages available on GitHub (for DunedinPACE, https://github.com/danbelsky/DunedinPACE; for the PC Clocks, https://github.com/MorganLevineLab/PC-Clocks).

### 4.2. UK Biobank Study

Using the interactive webtool developed by Deng et al 2024 (https://proteome-phenome-atlas.com/)(38), we extracted the associations between GDF15 and education, as well as GDF15 and mood disorder prevalence and incidence from the *UK Biobank (UKB)* dataset.

Briefly, in *UKB*, diseases were identified through linked electronic health records and classified using ICD-10 codes from hospital inpatient records, defining prevalent and incident diseases based on whether events occurred before or after participants’ baseline visits. Incident diseases were analyzed as time-to-event data, excluding participants with prior diagnoses, with follow-up ending at diagnosis, death, or the last record date (November 2023). Blood proteomic data were generated by the UKB-PPP consortium using Olink’s Proximity Extension Assay and sequencing to quantify 2,923 proteins across cardiometabolic, inflammation, neurology, and oncology panels. Following stringent quality control to address batch effects and technical variability, 2,920 proteins with ≤50% missing data were included in the analysis, with results expressed as Normalized Protein eXpression (NPX) values. In short, GDF15 was modeled as a continuous predictor using NPX values, which are log_₂_-transformed and normalized measures of relative protein abundance. A 1-unit increase in NPX corresponds to a doubling of GDF15 protein concentration; therefore, hazard ratios reflect the change in risk per doubling of GDF15 levels.

### 4.3 MiSBIE study

#### 4.3.1 Participants

Participants were enrolled in the *Mitochondrial Stress, Brain Imaging, and Epigenetics (MiSBIE) study*(65) in adherence to the directives outlined by the New York State Psychiatric Institute IRB protocol #7424, clinicaltrials.gov #NCT04831424. Recruitment was conducted both within our local clinic at the Columbia University Irving Medical Center, and nationally throughout the United States and Canada. All enrolled participants provided written informed consent, authorizing their participation in the investigative procedures and the potential dissemination of data. Recruitment occurred from June 2018 to May 2023 for a total of 70 healthy controls (females n=48, males n=22). This manuscript specifically reports on *MiSBIE* GDF15 dynamics in blood and saliva. The participants in this study were used as control group in a separate report on GDF15 dynamic in mitochondrial disease(62). Sex was determined by self-report. This single-step method is limited, does not distinguish between gender and sex, and can exclude both transgender and intersex people; current best practices include a two-step(103) measurement and broadening the available answer options.

Our inclusion criteria included English-speaking healthy adults willing to provide saliva samples and have blood collected using an intravenous catheter during the hospital visits, and the absence of pregnancy. Exclusion criteria included the presence of pronounced cognitive impairment precluding informed consent, recent occurrences of flu or other temporally pertinent infections within the four-week window preceding study participation, Raynaud’s syndrome, engagement in ongoing therapeutic or exercise trials registered on ClinicalTrials.gov, and the existence of metallic elements within or on the body, alongside claustrophobia posing an impediment to magnetic resonance imaging (MRI). In-person laboratory procedures were performed over 2-days (Tuesday and Wednesday), followed by a 1-week home-based saliva collection protocol.

Non-stress plasma and saliva samples were collected both in the morning, under fasting conditions around 10:00 AM, or after lunch around 1:30 PM. Both the breakfast and lunch meals were selected from the *MiSBIE* study menu to avoid large differences in meal types between participants.

#### 4.3.2 Stress reactivity

A modified Trier Stress Task (TSST) in the form of social-evaluative speech stress task was conducted on all *MiSBIE* participants on Day 1, as described previously(104, 105). The 5 min task included 2 min of preparation and 3 min of delivering a simulated speech to a court judge where participants defend themselves against an alleged shoplifting transgression. The speech was delivered in front of a confederate evaluator wearing a white lab coat (adding gravity to the situation), a mirror where participants can see themselves, and a (non-operational) video camera. To minimize participant discomfort during the task and avoid venipuncture-related stress responses during the task, an intravenous catheter was placed in the right arm >45 min before the initiation of the speech stress task and participants sat quietly, undisturbed for 30 minutes before receiving instructions for the speech task. Blood and saliva samples were collected at −5, +5, 10, 20, 30, 60, 90, and 120 min relative to the beginning of the social-evaluative stress task. Concurrently, during each blood collection, saliva samples were collected using salivettes (Starstedt Cat# 51.1534.500). This sampling strategy allows to capture both the reactivity and recovery phases of the stress responses. On day 2, while in the MRI scanner, to avoid habituation to the speech task, participants were informed that their performance on the previous day was “below average compared to other participants” and asked to prepare a speech to a court judge in defense of another alleged transgression (running a stop sign). Saliva samples were collected using salivettes at 3 timepoints: ∼30 min before the MRI session, 33 min into the session (∼9 min before beginning the speech task), and 75 min into the session (∼18 min after the beginning of the speech task).

#### 4.3.3 Plasma and saliva collection

The blood and saliva collection protocols are illustrated in Supplemental Figure S2A. Plasma was collected via a central venous catheter at 9 time points: 1) day 1 morning fasting sample, 2) 8 afternoon samples collected during the stress reactivity protocol. A total of 16-20mL of whole blood was collected at the morning fasting timepoint in two 10 mL K_2_EDTA blood collection tubes (BD #366643). A total of 5mL of whole blood was collected at each afternoon time point both before and after stress in 6mL K_2_EDTA blood collection tubes (BD #367899).

Saliva was collected at 26 timepoints: 1) day 1 morning fasting sample, 2) 8 afternoon samples collected during the stress reactivity protocol, 3) 1 sample after a cold pressor test, 4) day 2 morning fasting sample, 5) 3 afternoon stress samples before and during the MRI, 6) 12 at home samples collected across 3 days: (i) immediately upon awakening, (ii) 30 min after waking up, (iii) 45 min after waking up, and (iv) at bedtime. A total of 1-2 mL of saliva was collected at each sample timepoint using a salivette (Starstedt # 51.1534.500) following the Biomarker Network recommended procedure (106). Participants were directed to position the cotton swab at the center of their tongue within their mouth for 2 to 5 min. It was emphasized that they should refrain from biting down on the cotton and ensure the swab did not come into contact with their cheeks. Afterward, participants reinserted the cotton swab into the salivette recipient tube. Samples were collected on visit days 1 and 2, the salivettes were placed on ice (4°C) in a styrofoam box for transportation and further processing at the laboratory. For home-based sample collection, salivettes were frozen immediately after collection in a home freezer (typically −20°C). During the at-home morning collections, individuals were advised to ideally delay tooth brushing and eating until after the third sample (+45 min), while the nighttime sample was to be taken prior to bedtime tooth brushing. If participants needed to, they were advised to eat breakfast within the 30min break of awakening sampling and note the time and content. Participants were instructed to avoid consuming water or any other liquids within 10 minutes of each saliva sample collection. This was monitored and controlled for the laboratory samples. Within 2 weeks of collection, participants either transported (∼10%) or shipped them (∼90%) to the laboratory in a non-temperature controlled, pre-stamped USPS priority shipping.

#### 4.3.4 Biofluids (Plasma and Saliva) processing and storage

Whole blood tubes for morning fasting plasma samples were immediately inverted 10-12 times and centrifuged at 1000xg for 5 min at room temperature after collection. Samples were placed on ice (4°C) in a styrofoam box and transported to the laboratory for further processing. When samples reached the laboratory, they were immediately centrifuged at 2000xg for 10 min at 4°C. To avoid platelets and other cellular debris, approximately 80% of the plasma was collected from the upper portion of each tube, transferred and pooled into a fresh 15 mL conical tube. To further eliminate cellular components, pooled plasma tube was centrifuged at 2000xg for 10 min at 4°C. Around 90% of the resulting plasma supernatant was transferred to fresh 15 mL conical tube and mixed through inversion. The resulting cell-free plasma was aliquoted into 0.5-1.5 mL aliquots and stored at −80°C before being used for assays.

Stress reactivity time points include 8 tubes for each timepoint, starting at −5 min to 120 min. Whole blood tubes for afternoon stress plasma samples were immediately inverted 10-12 times and centrifuged at 2000xg for 3.5 min at room temperature after collection. Samples were placed on ice (4°C) in a styrofoam box and transported to the laboratory together for further processing. When samples reached the laboratory, they were immediately centrifuged at 2000xg for 10 min at 4°C. To avoid platelets and other cellular debris, approximately 80% of the plasma was collected from the upper portion of each tube and transferred into fresh 8*15 mL conical tubes. To further eliminate cellular components, plasma tubes were centrifuged at 2000xg for 10 min at 4°C. Around 90% of the resulting plasma supernatant was transferred to fresh 8*15 mL conical tubes and mixed through inversion. The resulting cell-free plasma was aliquoted into 0.5-1.5 mL aliquots for each timepoint and stored at −80°C before being used for assays.

When saliva samples reached the laboratory, Salivettes were centrifuged at 1000xg for 5 min in a refrigerated centrifuge at 4°C. To avoid cellular contamination with leukocytes or epithelial buccal cells(107), supernatant was carefully removed from the top of the tube, transferred to cryovials, and stored immediately at −80°C. Before use, saliva was thawed and centrifuged at 5000xg for 10 min to further eliminate cellular components. The resulting supernatant was collected as cell-free saliva and was used for subsequent assays.

### 4.4 The Mitochondria and Psychological Stress (MaPS) study

#### 4.4.1 Participants

Generally healthy community volunteers were recruited to participate in the *MaPS* study (clinicaltrials.gov #NCT04078035). Eligibility criteria were (1) 20-50 years of age (equal numbers within each decade of age); (2) for females, regular menstrual cycles over past 12 months, not lactating or pregnant; (3) generally physically healthy (no reported history of chronic systemic immune, metabolic or mitochondrial diseases, or chronic diseases that influence the central nervous, autonomic nervous or neuroendocrine systems); (4) not taking medications that influence the central nervous, autonomic nervous or neuroendocrine systems; (5) no reported history of psychotic illness or mood disorder; (6) resting blood pressure<140/90mmHg; (7) not a current smoker or user of illicit drugs; (8) weight>110lbs and BMI<30; and (9) fluent in English (used everyday in speaking and reading for at least 10 years). All enrolled participants provided written informed consent as approved by the University of Pittsburgh IRB. Recruitment occurred between July 2020-December 2022. A total of 72 participants were enrolled (n=34 self-identified as female; 38 as male). Participants were rescheduled if (1) they endorsed having an infection, cold or flu, taking antibiotics or glucocorticoids, or having received a vaccination or tattoo in the prior two weeks; (2) presented with symptoms in a range consistent with possible infection on the day of the visit; or (3) reported having taken glucocorticoids in the past 24 hours on the day of the visit.

#### 4.4.2 Stress Reactivity

Participants attended two afternoon laboratory sessions scheduled at least one month apart at the Behavioral Physiology Laboratory, University of Pittsburgh. Both sessions started between 1:00-3:00 PM. Participants were asked to avoid eating or drinking after 12 pm prior to each session. The order of sessions was counterbalanced in randomized starting order across participants. At one session, they completed the same 5-minute social-evaluative speech task that was used in *MiSBIE* (see above). At the other session they rested quietly for the 5-minute task period and were not exposed to the stressor. On both occasions, an intravenous catheter was inserted and participants sat quietly for a 30-minute baseline (habituation) period before completing the task. Following the task period, participants watched a wildlife documentary for a 120 min recovery period. Participants were left alone as much as possible, except when the study nurse had to re-insert or reposition the catheter, during all periods except when performing the speech task. Blood samples were collected 5 min before and 5, 10, 20, 30, 45, 60, 75, 90, and 120min after the beginning of the task period. Saliva was not collected to avoid potential interference with the participant’s psychological state. Blood was drawn without the participant’s awareness via an obscured IV line running through the wall behind the participant. This protocol variation minimizes psychological interference after the induction of the stressor. However, it also eliminated interpersonal interactions for 2 hours after the stressor, which could produce anxiety.

### 4.5 Intensive repeated-measures n=2 study

#### 4.5.1 Participants and procedures

To evaluate the intra-individual fluctuations of salivary GDF15, repeated daily measurements were taken over periods of 53 and 60 days from two white male participants: author M.P., 34 years old (Participant A) and author G.G., 35 years old (Participant B, see(108)). Study investigators served as participants to mitigate the excessive demand on volunteer participants required to collect numerous daily samples over multiple consecutive weeks. The study was approved by the New York State Psychiatric Institute (Protocol #7748). Participants provided written informed consent for participating and publishing the results. As for the at-home collection protocol in the *MiSBIE* cohort, participants collected saliva immediately upon awakening, 30min and 45min after waking up, and at bedtime. Participants delayed consuming liquid and tooth brushing until after the third sample (45 min), and the nighttime sample was taken prior to bedtime tooth brushing. On each morning and evening, participants completed a morning diary assessing positive and negative affective states, using items adapted from the modified Differential Emotional Scale (MDES), including ratings of stress levels on a 5-point Likert scale: 1(Not at all), 2 (A little bit), 3 (Somewhat), 4 (Moderately) to 5 (Extremely)(109).

#### 4.5.2 Saliva processing and storage

Salivette samples were stored in home freezer (−20°C) before being transported or shipped frozen to the laboratory. Upon arrival salivettes were thawed at room temperature and centrifuged at 1000 x g for 5 min in a refrigerated centrifuge operating at 4°C to extract saliva from the cotton swab. 200uL of saliva supernatant was taken from the top, transferred to a 1.5 ml Eppendorf tube and stored at −80°C. Prior to analysis, Eppendorf tubes were further centrifuged at 2000 x g for 10 min at 4°C to remove cellular debris, and the resulting supernatant was collected as cell-free saliva and was used for GDF15 measurements by ELISA. To control for the effect of saliva protein concentration on GDF15 results, total protein concentration was measured using the Bicinchoninic acid (BCA) assay as described previously(108).

### 4.6. In house GDF15 assay

Plasma, serum and saliva GDF15 levels collected in *MiSBIE* study, *MaPS* study, *Intensive repeated-measures n=2* study, and *Overnight meal alignment study* were quantified using a high-sensitivity ELISA kit (R&D Systems, DGD150, SGD150) following the manufacturer’s instructions. See Dataset S1 for raw GDF15 data from each study. Different lot numbers were used, and the average coefficient of variation (C.V.) between lot numbers was determined by reference samples for quality control. Plasma and serum samples were diluted with assay diluent (1:4 ratio) to maximize the number of samples within the dynamic range of the assay. Saliva samples were not diluted. Absorbance was gauged at 450nm, and concentrations were computed utilizing the Four Parameter Logistic Curve (4PL) model.

Samples were run in duplicates, on separate plates when possible, and the concentration for each sample was computed from the average of the duplicates. Standard curve (5 samples per plate) and plasma reference samples (2-3 samples per plate, same sample per batch) were run with each individual assay and the inter-assay C.V. was monitored. All standard curves and references were overlaid on top of each other to monitor failed runs. Samples were run in duplicate plates when possible and those with C.V. larger than 15% were rerun. When it was not possible to rerun (e.g., no sample left), sample sets with a C.V. >15% between the duplicates were included. We performed sensitivity analyses excluding these samples, confirming that the results were unchanged by the presence of absence of these samples with lower reliability. Values below the mean minimum detectable dose (2.0pg/ml) were considered as non-detectable (reported as NA) and excluded in the graphs or statistical analyses. For the *MiSBIE*, samples were run in 3 batches over 2 years, multiple quality control measures were applied to monitor batch-to-batch and within batch variability. Data-preprocessing and quality control measures was done using R Software (version 4.2.2). For the number of samples processed in-house, broken down by study and sample type, along with the proportion of missing data, see Dataset S2.

Based on the 1000 Genomes Project, around 15-30% of the population assessed in the project carried the common H202D variant in GDF15(94). Compared to the Roche Elecsys assay, the R&D ELISA kits was found to report GDF15 concentrations that are 36% lower in individuals carrying one D allele or 61% lower in those that carry two D alleles at position 202 of the pro-peptide (or position 6 of the mature peptide)(95). Despite this limitation, our within-person findings related to psychosocial stress reactivity, diurnal variation, and awakening responses would be unaffected by the underestimation of GDF15 concentration in some participants with the H202D variant.

### 4.7. Statistical analyses

For *MiSBIE*, *MaPS*, and *Intensive repeated-measures n=2 study*, within individuals comparisons (e.g., morning to afternoon, fasting day 1 to day 2, −5 vs. 5 min, participant A vs B) were assessed using the paired Wilcoxon signed-rank test, while between-individual comparisons (e.g., fasting morning male vs. female) were assessed using the Mann-Whitney test. Stress reactivities were quantified as the percentage change from baseline. Mixed-effects models were used to evaluate the effect of acute stress or diurnal response on respective biomarker over time and Tukey’s multiple comparison test was applied for post-hoc analyses.

Two-way ANOVA was used to test the effects of acute stress or diurnal response on respective biomarker overtime, group (e.g. *MiSBIE* male vs female, *MaPS* control vs stress), and their interaction, followed by Tukey’s multiple comparison test.

Associations between continuous variables (e.g., GDF15 vs. catecholamine reactivity, GDF15 vs. saliva cortisol reactivity, etc.) were analyzed using Spearman rank correlation. In *MiSBIE*, plasma GDF15 stress reactivity was calculated as the fold-change from baseline (−5 min) to the peak level at or before 10 minutes post-stress, with reactivity gated at 10 minutes based on the group-average peak. Saliva GDF15 stress reactivity was calculated as the fold-change from baseline to the group-average peak at 5 minutes. Cortisol stress reactivity was calculated as the fold-change from baseline (−5min) to the peak level at or before 30 minutes post-stress. Norepinephrine and epinephrine stress reactivity was calculated as the fold-change from baseline −5min) to the peak level at or before 20 minutes post-stress. Plasma lactate reactivity was calculated as the fold-change from baseline to the peak at 5 minutes, the group-average peak timepoint. In *MaPS*, plasma GDF15 reactivity was calculated as the fold-change from baseline (−5min) to the group-average peak at 5 minutes. Cortisol stress reactivity was calculated as the fold-change from baseline (−5min) to the peak level at or before 30 minutes post-stress. Norepinephrine and epinephrine stress reactivity was calculated as the fold-change from baseline (−5min) to the peak level at or before 20 minutes post-stress. In *MiSBIE* and *Intensive repeated-measures n=2 study*, the magnitude of the GDF15 awakening response was calculated by subtracting the 45min level from the peak morning value (across awakening, 30, and 45min), the magnitude of the cortisol awakening response was calculated by subtracting the awakening level from the peak morning value (across awakening, 30, and 45min).

For *FHS*, GDF15 was natural log-transformed to normalize skewed distributions and standardized. Multivariable linear regression was conducted with GDF15 as dependent variables, chronic-stress related factors as independent variables, and chronological age and sex as covariates. No adjustments were made for multiple testing due to the hypothesis-generating nature of the study. For epigenetic clocks regression analyses, methylation measures were adjusted for batch effects by regressing the measures on batch variables and using the residuals. Since plasma GDF15 levels significantly correlate with age (r=0.60, p<0.0001) and is higher in men compared to women (p=0.0059), all *FHS* analysis, unless otherwise noted, were performed on age- and sex-corrected GDF15 to ensure that observed associations are independent of these confounding variables. Age and sex were also regressed out from each epigenetic clock to ensure comparability with GDF15, unless otherwise noted. To account for potential confounding from variations in immune cell types, relative abundances of 12 immune cell subtypes was estimated using established R package and sensitivity analysis was performed. We report effect sizes in terms of GDF15 standard deviation units in the FHS; Hedges’ g was used to allow unbiased comparison across analyses with varying sample sizes.

All data-preprocessing and quality control measures was done using R Software (version 4.2.2 and 4.3.0) and are available as supplemental files. Statistical analyses were conducted using R (version 4.2.2 and 4.3.0) and GraphPad Prism (version 9.4.1).

## 5. Code and data availability statement

Data from *FHS* was accessed from the NIH database of Genotypes and Phenotypes (dbGaP; accession number phs000724.v9.p13). Data from *UKB* was generated using the proteomic atlas(38) (https://proteome-phenome-atlas.com). Data from *MiSBIE* study, *MaPS* study, *Intensive repeated-measures n=2* study, and *Overnight meal alignment study* is available upon request. The code use for data analysis in this manuscript is available on GitHub (https://github.com/mitopsychobio/2024_GDF15_Dynamics_Huang).

## Author contributions

C.T., D.W.B. and M.P. conceived and supervised the overall project. M.P., M.H., C.T., R.P.S., V.L., R.P.J. developed *MiSBIE*. A.L.M., B.A.K., M.P. developed *MaPS*. M.P., G.G., S.R., C.T. developed *Intensive repeated-measures n=2 study*. M.P.S.O. developed *Overnight Meal Alignment Study.* C.E.I, M.J.S. and D.W.B. curated GDF15 and epigenetic clocks data in FHS. C.K. and K.E. recruited *MiSBIE* participants. C.K. and S.R. enrolled participants, collected study visits data and samples. M.K. processed samples. S.L. and M.H. performed the medical examinations. Q.H., R.H., S.R., A.S.M., C.C.L. performed GDF15 assays. E.T. performed the lactate assays in collaboration with J.M. Q.H., A.S.M., C.C.L., J.D., N.B.A. performed statistical analyses. Q.H. and C.T. prepared the figures. D.W.B. advised on statistical analyses. C.T., Q.H., M.P. and D.W.B. drafted the manuscript. All authors reviewed the final version of the manuscript.

## Acknowledgements

Work of the authors is supported by R01MH122706, R01AG066828 and Baszucki Brain Research Fund to M.P., the Wharton Fund to M.P. and C.T., R01MH119336 to A.L.M., B.A.K., and M.P., R01HL142648, R35HL155670 and UL1TR001873 to M.P.S.O., R01AG073402 to D.W.B. (a fellow of the CIFAR CBD Network), the New York Obesity Nutrition Research Center grant P30DK026687-39, the Chercheur-boursier Junior 2 salary award from the Fonds de recherche du Québec en santé (FRQS-297877) to G.G., the Chercheur-boursier Junior 2 FRQS (332273: https://doi.org/10.69777/332273) and the Fondation de l’Institut universitaire en santé mentale de Montréal to R.P.J. The Framingham Heart Study is conducted and supported by the National Heart, Lung, and Blood Institute (NHLBI) in collaboration with Boston University (Contract No. N01-HC-25195, HHSN268201500001I and 75N92019D00031). This manuscript was not prepared in collaboration with investigators of the Framingham Heart Study and does not necessarily reflect the opinions or views of the Framingham Heart Study, Boston University, or NHLBI. The DNA methylation data analyzed were supported by funding from the NIH including funds from the NHLBI, Division of Intramural Research (D. Levy, PI) and the NIH Director’s Challenge Award (D. Levy, PI)

