## Supplemental Figures 1-6 for "The energetic stress cytokine GDF15 is elevated in the context of chronic and acute psychosocial stress"

### 1. Supplemental Figures

#### Supplemental Figure 1

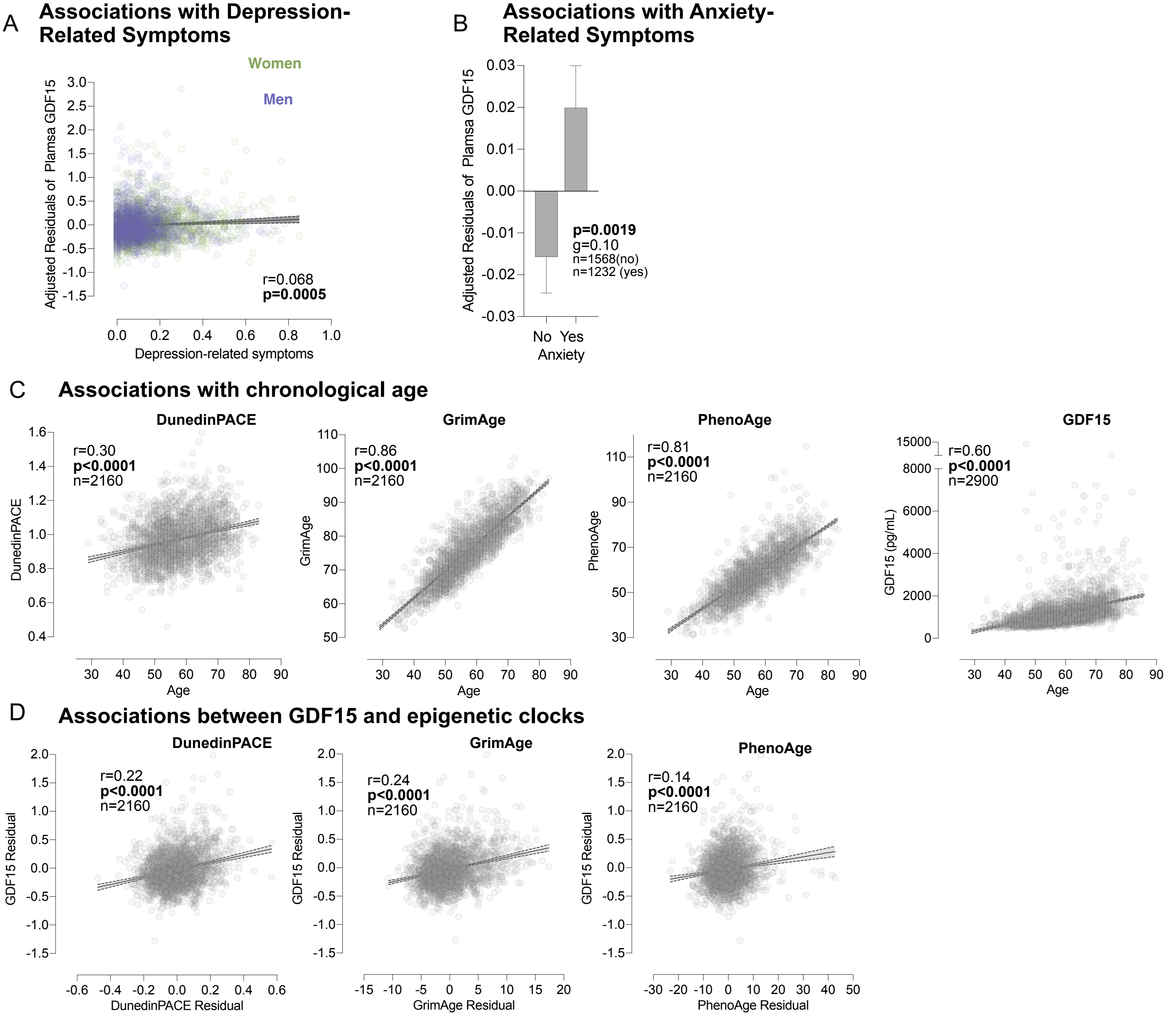


**Figure S1. Associations of GDF15 with depression and anxiety symptoms, chronological age, and epigenetic clocks in FHS.**

(**A**) Scatterplot of the association between age-corrected GDF15 levels and depression-related symptoms. (**B**) Average age-corrected GDF15 levels by anxiety-related symptoms in (n=1572 for 'No', n=1232 for 'Yes'). (**C**) Scatterplot of the associations between associations between age and GDF15 (n=2,900) and epigenetic clocks (n=2,160). (**D**) Scatterplot of the associations between age-corrected plasma GDF15 and each epigenetic clock (DunedinPACE, GrimAge, PhenoAge).
P-values and effect sizes from (A,C,D) Spearman's rank correlations and (B) Mann-Whitney rank t-test and Hedge's g. (B) Data shown as mean ± SEM. Age corrected GDF15 and epigenetic clocks values were used for all analysis and graphing.

#### Supplemental Figure 2

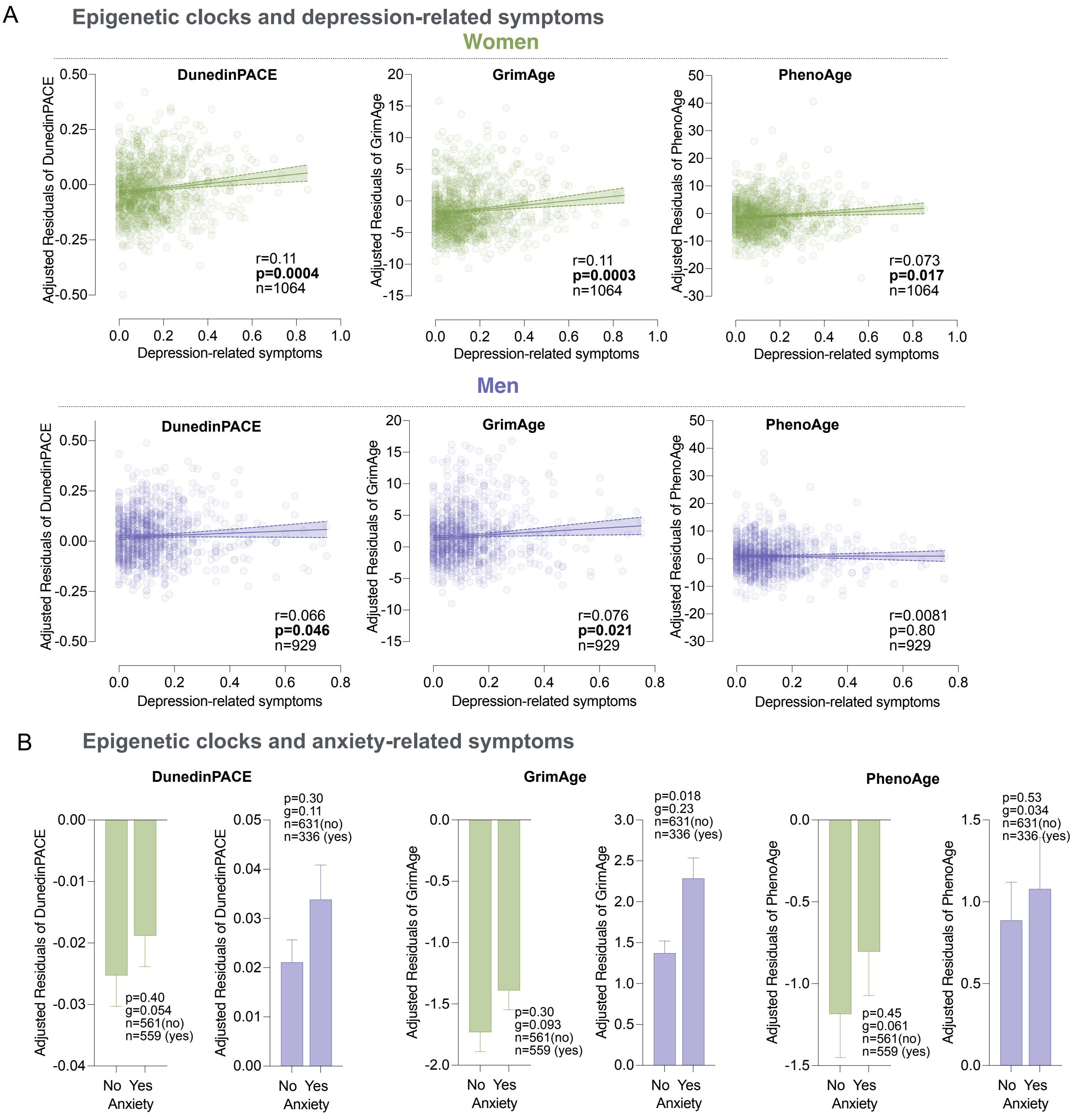


**Figure S2. Associations of epigenetic clocks with depression- and anxiety-related symptoms in FHS.**

(**A**) Scatterplot of the associations between epigenetic clocks and depression-related symptoms, presented separately for women and men. (**B**) Average epigenetic clocks by anxiety-related symptoms, presented separately for women (left) and men (right).
P-values and effect sizes from (A) Spearman's rank correlations and (B) Mann-Whitney rank t-test and Hedge's g. (B) Data shown as mean ± SEM. GDF15 and epigenetic clocks values used in analysis and graphing were age corrected.

**Supplemental Figure 3**

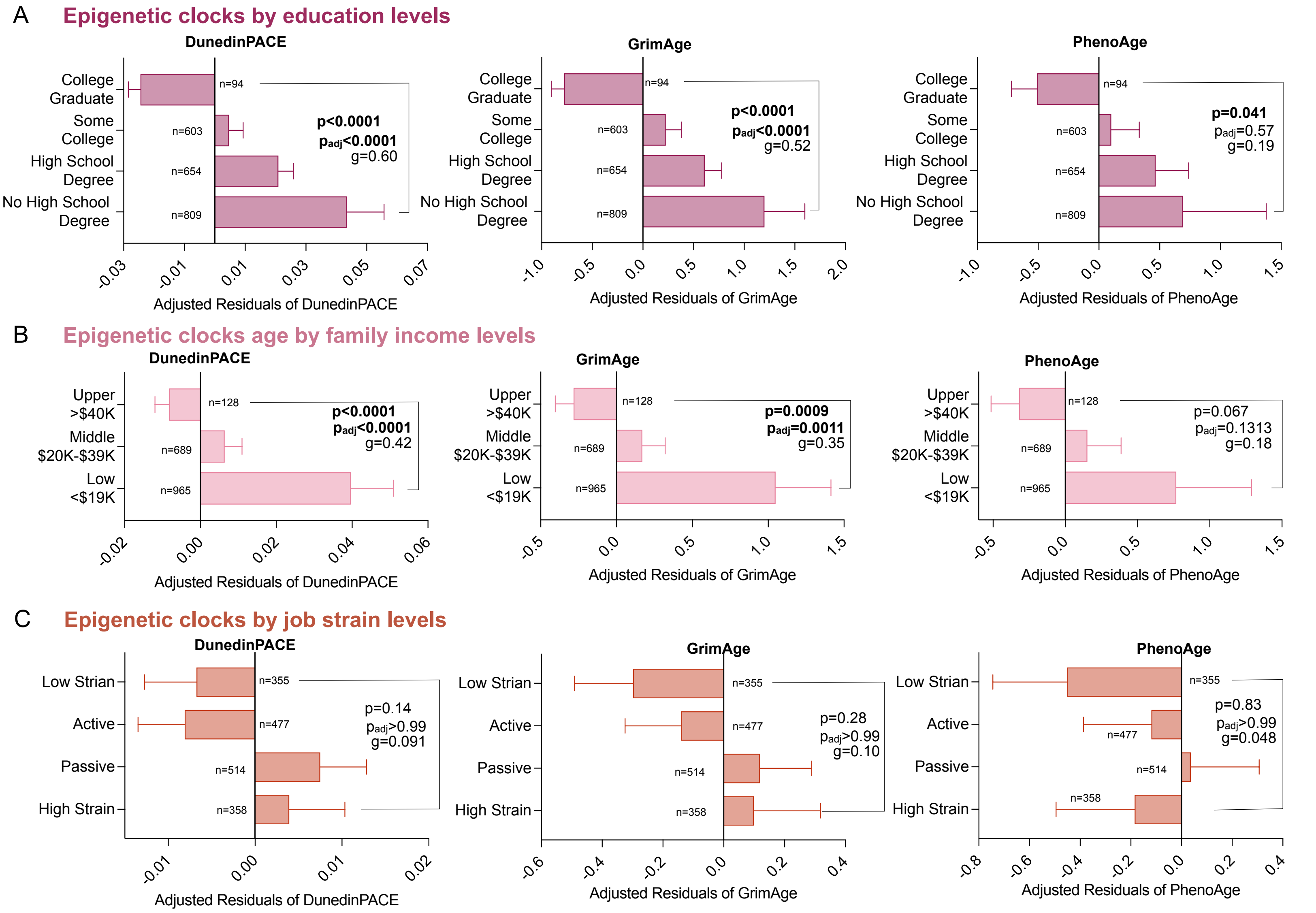


**Figure S3. Associations of plasma GDF15 and epigenetic clocks with socioeconomic and aging factors in FHS**

Average epigenetic clock levels (n=2160) by (**A**) education levels (n=2160), (**B**) family income levels (n=1708), (**C**) job strain (n=1704) defined as 4 categories: low Strain (high in decision latitude and low on psychological job demand); active (high on both decision latitude and psychological job demand); passive (low on both decisions attitude and psychological job demand); high strain (low on decision latitude and high on psychological job demand). P-values from Krusakal-Wallis test and Dunn's multiple comparison test adjusted for multiple testing, data shown as mean ± SEM. Effect sizes estimated with Hedges' g. GDF15 and epigenetic clocks values used in analysis and graphing were age and sex corrected. *p<0.05, **p<0.01, ***p<0.001, ****p<0.0001.

#### Supplemental Figure 4

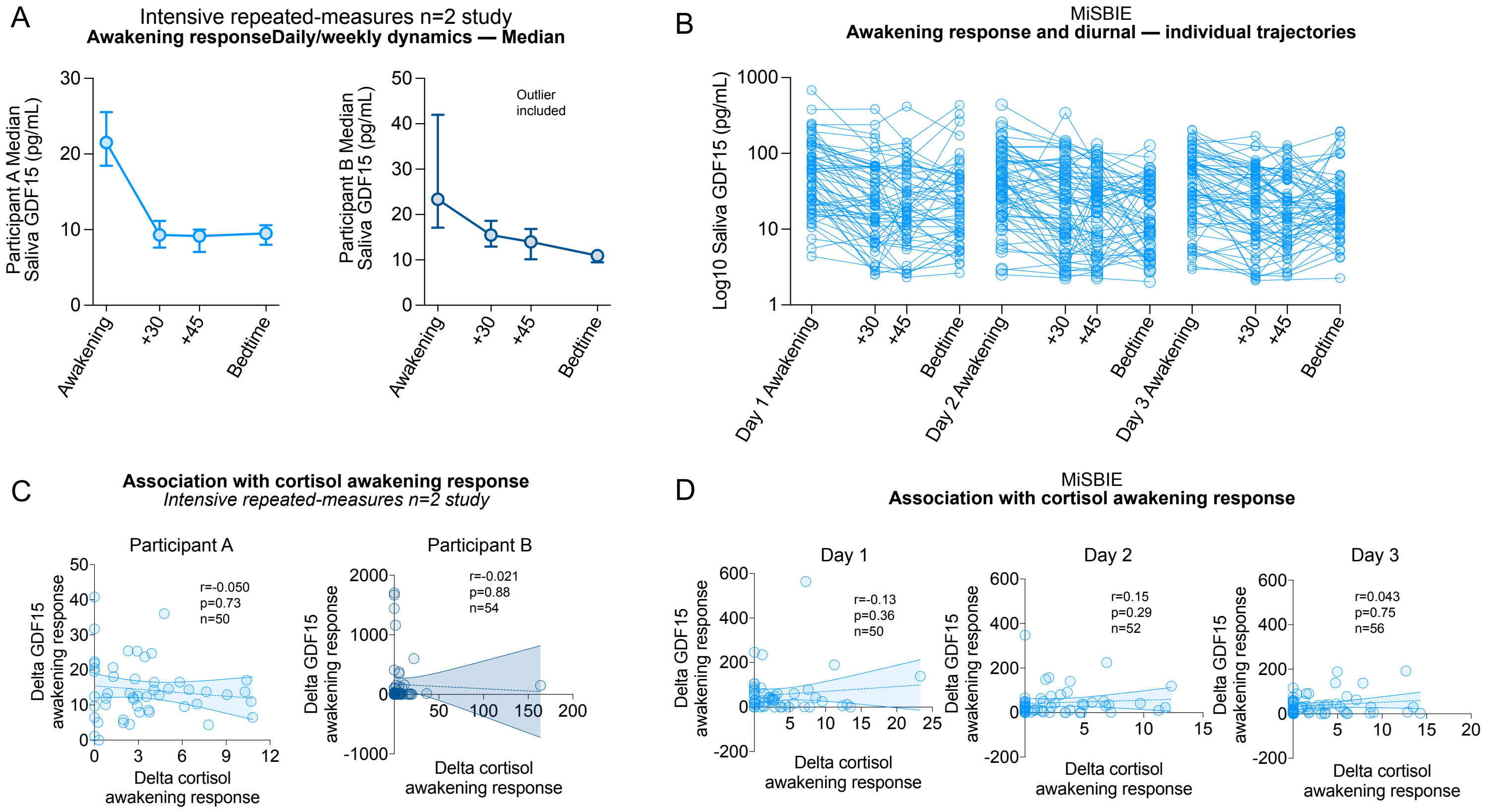


**Figure S4. Awakening dynamic behavior and sex comparison of GDF15 in human saliva.**
(**A**)Intensive repeated-measures n=2 study 53-day (Participant A, left) and 60-day (Participant B, right) saliva awakening GDF15 profiles group shown as median. (**B**) Individual trajectories of saliva GDF15 (value graphed as log10) diurnal variation from MiSBIE healthy controls (n=678 observations, 67% female) on three separate days at awakening, 30 and 45 min after awakening, and bedtime. (**C**) Correlation between cortisol awakening response and GDF15 awakening response in Participant A and Participant B across 53 days and 60 days, respectively. Each dot represent an association from a day. (**D**) Correlation between cortisol awakening response and GDF15 awakening response in MiSBIE. Correlations between cortisol and GDF15 awakening response was analyzed separately for each day. Each dot represent an association from a participant in a day.
(B) Data shown as median ± 95% CI. (C,D) The delta of GDF15 awakening response was calculated by subtracting the 45min level from the peak morning value (across awakening, 30, and 45min), the delta of cortisol awakening response was calculated by subtracting the awakening level from the peak morning value (across awakening, 30, and 45min). P-values from Spearman's rank correlations.

**Supplemental Figure 5**

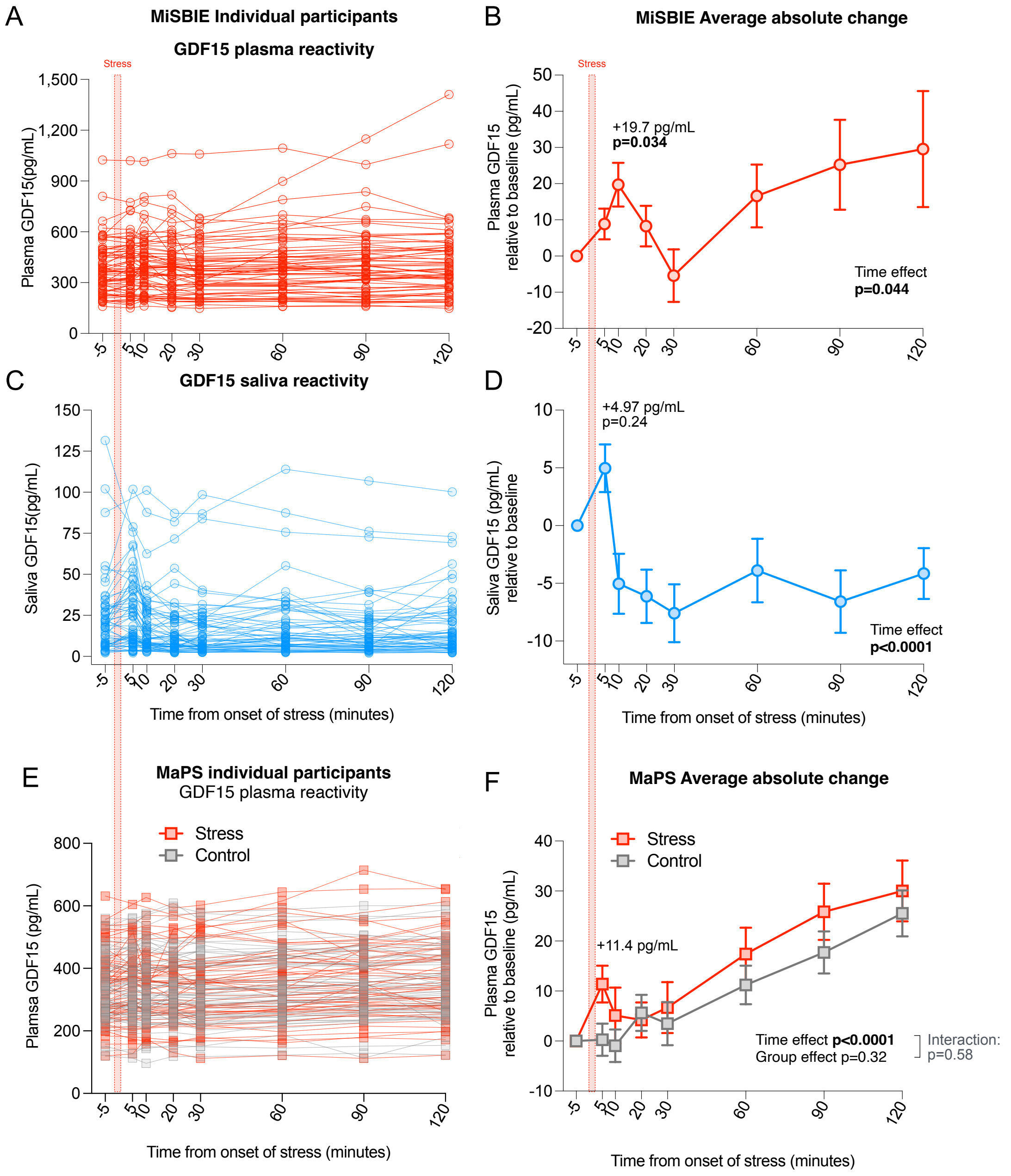


**Figure S5. Social-evaluative stress acutely increases plasma and saliva GDF15**

(**A,B**) MiSBIE plasma GDF15 levels before and after psychological stress shown as (**A**) individual trajectories and (**B**) absolute change from baseline (-5 min). n=65, 68% female, 512 observations.(**C,D**) Same as in (**A,B**) for Saliva GDF15. n=57, 65% female, 442 observations.(**E,F**) MaPS Plasma Plasma GDF 15 levels for control and stress visit shown (**E**) individual trajectories and (**F**) absolute change from baseline (-5min). n=70, 48% female, 916 observations.
Data shown as mean ± SEM. P-values from (B,D) mixed-effect model followed by Tukey multiple comparison test corrected for multiple testing and (F) two-way ANOVA followed by Tukey multiple comparison test, corrected for multiple testing. *p<0.05, **p<0.01, ***p<0.001, ****p<0.0001.

**Supplemental Figure 6**

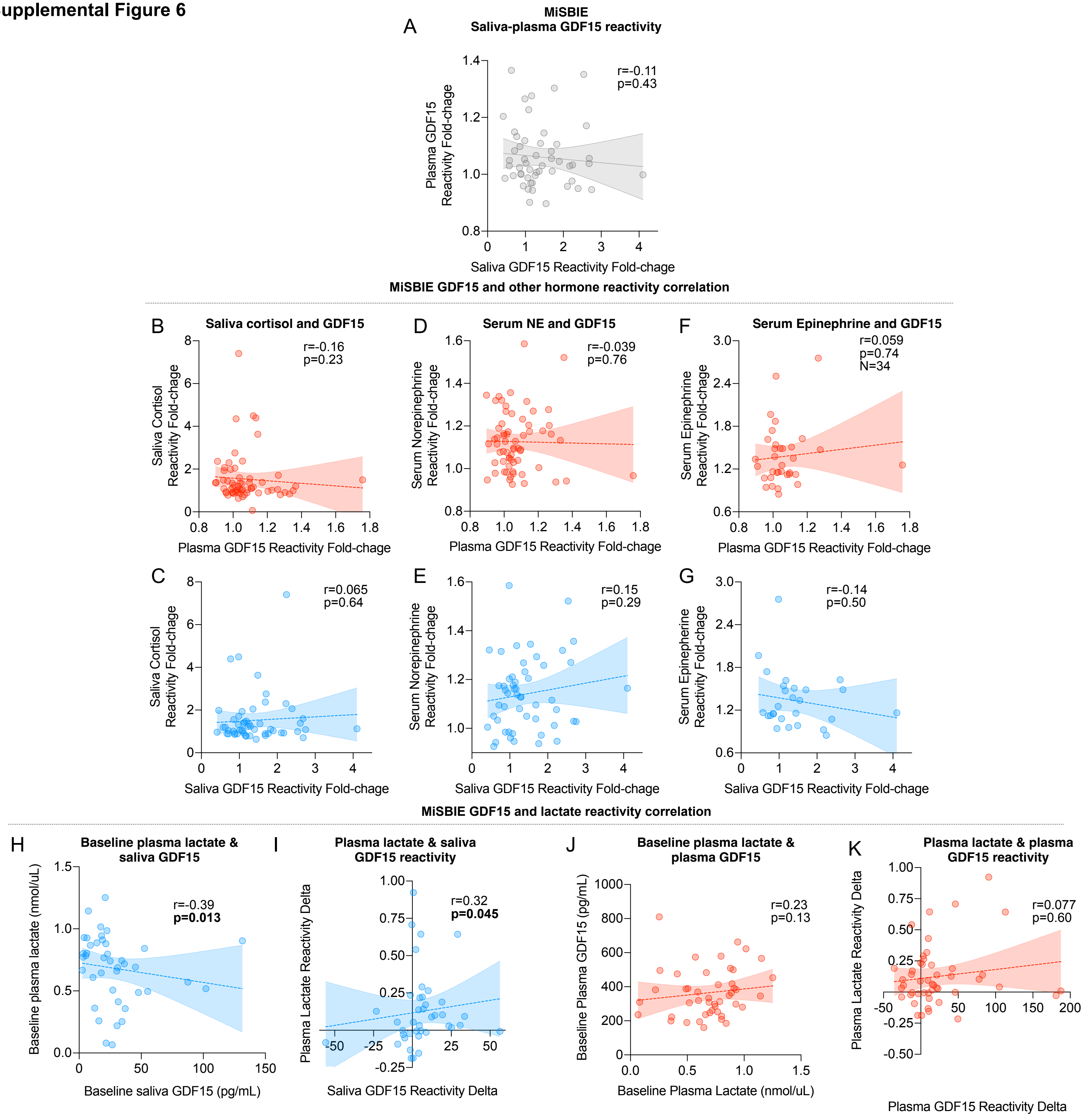


**Figure S6. Associations of plasma and saliva GDF15 stress responses with cortisol, catecholamines, and lactate in MiSBIE.**
(**A**) Correlation between plasma and saliva GDF15 stress response measured in the same person (N=55, 69% female). Plasma GDF15 stress reactivity was calculated as the fold-change from baseline (-5 min) to the peak level at or before 10 minutes post-stress, with reactivity gated at 10 minutes based on the group-average peak. Saliva GDF15 reactivity was calculated as the fold-change from baseline to the group-average peak at 5 minutes.**(B,C)** Correlation between saliva cortisol response with **(B)** plasma (N=62 , 68% female) and (**C**) saliva GDF15 responses (N= 53, 68% female).Cortisol stress reactivity was calculated as the fold-change from baseline (-5 min) to the peak level at or before 30 minutes post-stress. **(D,E)** Correlation between serum norepinephrine response with **(D)** plasma (N=65, 68% female) and **(E)** saliva (N=52, 68% female) GDF15 responses. Norepinephrine stress reactivity was calculated as the fold-change from baseline (-5 min) to the peak level at or before 20 minutes post-stress. **(F,G)** Correlation between serum epinephrine response with **(F)** plasma (N=34, 75% female) and **(G)** saliva (N=27, 80% female) GDF15 responses. Epinephrine stress reactivity was calculated as the fold-change from baseline (-5 min) to the peak level at or before 20 minutes post-stress**(H,I)** Association between (**H**) baseline levels (N=41) and (**I**) stress response (N=40) of plasma lactate and saliva GDF15. Plasma lactate reactivity was calculated as the fold-change from baseline to the peak at 5 minutes, the group-derived peak timepoint. **(D,E)** Same as in **(B,C)** for plasma GDF15. Stress response N=48, baseline levels N=46. P-values from Spearman's rank correlation. *p<0.05, **p<0.01, ***p<0.001, ****p<0.0001.
