## Supplemental Information Text for "The energetic stress cytokine GDF15 is elevated in the context of chronic and acute psychosocial stress"

### 2. Supplemental Information Text

#### *2.1 Day-night variation in blood GDF15*

To explore if the there is a day-night variation in blood GDF15, we analyzed samples collected in *the Overnight Meal Alignment Study* (See Supplemental Information Methods  below for details). Serum samples were collected via indwelling catheter and intravenous (IV) line every 30 minutes overnight during sleep, and in the morning after waking up, on two separate visits (n=6 participants). Some participants exhibited up to 25.6% variation in serum GDF15 levels between day and nighttime, consistent with a previous report(1). However, in this small sample, we found no systematic serum GDF15 diurnal variation across participants (SI Text Figure 1). Thus, blood GDF15 diurnal variation was either too small in magnitude or too idiosyncratic in its timing to be detected in this sample. The absence of a detectable pattern may also result from the limited sample size, and larger studies are needed to better characterize the diurnal regulation of blood GDF15

#### *2.2 Intensive daily repeated measures in n=2.*

In *Intensive repeated-measures n=2 study*, we performed an intensive repeated-sampling protocol for saliva on two male participants using the same 4-timepoints as in *MiSBIE*. Both participants collected saliva for 53 (Participant A) and 60 (Participant B) consecutive days, as described previously(2). This yielded 199 and 221 GDF15 measures for participant A and B, respectively.

*Participant A* (male, age 34, sedentary, stress score average: 0.90 out of 5 on the modified Differential Emotional Scale (mDES)(3)) saliva GDF15 levels ranged from 2.3pg/mL to 54.3pg/mL (median: 10.5pg/ml, SI Text Figure 2A). Saliva GDF15 exhibited an average -57.1% decline from awakening to 30 min, and -60.5% at 45 min. At bedtime, saliva GDF15 levels were, on average -55.8% lower than at awakening, consistent with group results in *MiSBIE* (Figure 3B and 3D).

*Participant B* (male, age 35, competitive runner, stress score average: 2.57 out of 5 on the mDES) saliva GDF15 levels ranged from 3.5pg/mL to 4,428pg/mL (median: 15.1pg/ml, SI Text Figure 2A). His awakening response was on average -85.5% at 30 min, -88.7% at 45 min. At bedtime, saliva GDF15 levels were, on average -69.6% lower than at awakening (Figure 3C). During the data collection period, Participant B was actively training for a marathon (race on day 27, red arrow), possibly accounting for the high GDF15 values and the overall higher mean levels than Participant A. The highest GDF15 value (4,428pg/mL) was recorded at the 45 min timepoint the day after the marathon (day 28), consistent with prior reports linking exhaustive exercise with *blood* GDF15(4), but here observed punctually in *saliva*, at a specific timepoint, *the day after* the thorough effort (Figure 3E).

Within each participant, the three morning time points (awakening, 30, 45 min) were significantly correlated with one another (r=0.36-0.60, all p<0.01, SI Text Figures 2B-C). This result suggest robust day-to-day variation, marked by some low GDF15 mornings and some high GDF15 mornings. Morning and evening timepoints were not correlated, possibly pointing to different drivers of saliva GDF15 levels between morning and evening.

Cortisol levels were measured from the same saliva sample as GDF15. As reported in a previous study, Participant A exhibited positive cortisol awakening response in 80% of the 53 days while Participant B exhibited positive cortisol awakening response in 66% of the 60 days(2). No association was observed between the magnitude of the cortisol and GDF15 awakening responses in either participant (SI Text Figures 2D-E).

#### *2.3 Psychobiological regulation of plasma GDF15 in MaPS*

To validate the stress-induced changes in plasma GDF15 found in *MiSBIE*, we analyzed samples from the *MaPS* study – (*Mitochondria and Psychological Stress Study*, see Methods and Materials for details). In *MaPS*, 68 healthy individuals (mean age 30.6 years, 51.2% female) completed two laboratory visits in randomized order, involving either the same speech task as in *MiSBIE* or a control condition (no stressor beyond the blood draw and laboratory setting) (Figure 1D). Plasma GDF15 was measured at the same 8 timepoints as in *MiSBIE* (5 min before and 5, 10, 20, 30, 60, 90, 120 min after the initiation of the stressor). Acute psychosocial stress increased plasma GDF15 by 3.5% ([95%CI 1.4%-5.7%], p=0.0085) at the 5 min, validating the direction and magnitude of the *MiSBIE* results (Figures 1E, see Figure S4E-F for individual trajectory and average absolute change). On both control and speech task visits in *MaPS*, GDF15 gradually increased over the 2-hour post-stressor recovery period, reaching a maximum at 120min (+9.0% in the stress visit, +8.1% in the control visit, relative to baseline). Combined with our *MiSBIE* results, these findings suggest that psychosocial stress increases human plasma GDF15 levels within minutes.

In *MaPS*, participants randomized to the stress condition on their first visit did not significantly differ on their baseline plasma GDF15 concentrations compared those randomized to control on their first visit (p=0.45, SI Text Figures 3A-C). Interestingly, and consistent with this hypothesis and the aversive (e.g., nausea) experience of GDF15(5, 6), across the cohort, the two participants who did not return for their second visits (lost to follow-up) presented with the highest baseline GDF15 levels, which remained elevated throughout the stress task (SI Text Figures 3D-F).

In *MaPS*, levels of cortisol, norepinephrine, and epinephrine were assessed at 6 timepoints during the stress task (5 min before and 5, 10, 20, 30, 45 min after the initiation of the stressor). Consistent with our MiSBIE study findings (Figures S6B-G), we did not observe an association between the magnitude of the plasma GDF15 response and the response of each of the neuroendocrine factors (SI Text Figures 3G-I).

#### *2.4 Blood lactate stress reactivity and relationship with GDF15*

Using a subset of serum samples from *MiSBIE* (See Supplemental Information Methods below for details), we confirmed that, similar to what was previously observed in mice(7), evoked mental stress elevated circulating lactate in humans. Lactate is produced by increasing the intracellular NADH/$NAD^+$ ratio (i.e., reductive stress), which pushes the lactate dehydrogenase reaction towards lactate production(8). In *MiSBIE*, the brief mental stress increased plasma lactate by an average of 14.5% (p=0.0054, range: -39 to +151%, SI Text Figure 4A) from baseline to 5 min, demonstrating in humans that a brief mental stressor induces systemic reductive stress (see SI Text Figure 4B for individual trajectory and SI Text Figure 4C for absolute change).

Resting blood lactate levels were found to be negatively correlated with resting GDF15 saliva levels (r=-0.39, p=0.013; Figure S5H). However, resting blood lactate levels explained no more than 5-15% of the variance in inter-individual differences in resting plasma GDF15 or plasma GDF15 reactivity (Figures S5J-K).

#### *2.5 AM-PM dynamic variation*

To quantify and compare how much saliva and plasma GDF15 levels change over time, in *MiSBIE* we implemented a repeated-sampling design including 14 saliva collections over a 2-day period (SI Text Figure 5A). Frequent saliva sampling allowed for a refined description of GDF15 dynamics. We found that compared to plasma GDF15 concentrations, which have been shown to exhibit a moderate degree of stability over weeks(9), saliva GDF15 levels presented 10.0-fold higher within-person variations (SI Text Figures 5B-C). Thus, saliva GDF15 is approximately an order of magnitude more dynamic or changeable than plasma GDF15.

We then examined how much plasma and saliva GDF15 levels vary across consecutive days, and from morning to afternoon within a given day. In *MiSBIE*, morning fasting saliva collected under the same conditions on two subsequent days exhibited higher GDF15 levels on the first study visit day (when the hospital environment was novel) and decreased on average by 26% from Day 1 to Day 2 (p=0.032, SI Text Figure 5D), demonstrating significant within-person variation. This decline may also reflect possible habituation to the novel, and therefore psychologically stressful research environment.

We also had the opportunity to investigate changes in GDF15 from morning (9am, fasting) to afternoon (~1-2pm, fed) on the two consecutive days in *MiSBIE*. On Day 1, from morning to afternoon, GDF15 levels decreased on average by -12% (p<0.0001) in plasma but tended to increase by +6.5% (p=0.06) in saliva (SI Text Figure 5E). On Day 2 (only saliva collected), saliva GDF15 levels again increased from morning to afternoon by an average of +20% (p<0.0001). The magnitude and direction of change in saliva GDF15 between days 1 and 2 were not significantly correlated (r=0.05, p=0.76, SI Text Figure 5F). Thus, both the fasting/fed state(10) and the time of day might influence blood and saliva levels differently, further highlighting the distinct regulation of saliva GDF15 from circulating blood levels.

#### *2.6 Sex differences in saliva GDF15 awakening response in MiSBIE*

Building on the dynamic AM–PM variation in GDF15, we next examined whether sex differences were evident in the saliva GDF15 awakening response in the *MiSBIE* sample. Over the awakening time course, males tended to exhibit a 39-101% higher median saliva GDF15 levels than females (n.s.), although both exhibited a similar decline by 47-57% after waking up (SI Text Figures 6A-B). Thus, in the *MiSBIE* sample, saliva GDF15 exhibits rapid and robust variation after waking up, with potential sexual dimorphism in the absolute saliva GDF15 levels.

### 4. Supplemental Information Text Methods

#### *4.1 Overnight meal alignment study*

*Participants and procedures*

Healthy females and males were recruited to participate in a randomized crossover study testing the impact of varying mealtimes, relative to wake time, on energy balance regulation (clinicaltrials.gov #NCT03663530). Participants interested in the study underwent a multi-stage screening process. Inclusion criteria included being 20-49 years of age at the time of screening, having a body mass index between 20-34.9 kg/m$^2$, healthy habitual sleep patterns, and habitually consuming the first meal of the day within 1 h of waking. Individuals who were pregnant or lactating, smoked (currently or <3 years since stopping), reported excessive caffeine or alcohol intake, had extreme morning or evening chronotype, or who were actively engaging in intentional weight change or had a recent weight change >5% in the past 3 months were excluded. In addition, prospective participants with cancer; chronic cardiovascular, metabolic, autoimmune, neurologic, psychiatric, or sleep-related disorders/conditions; or who or take medications or supplements known to affect sleep or energy balance were not included cancer were not included. Individuals meeting all eligibility criteria competed a two-week outpatient screening period, which included daily assessment of sleep via wrist accelerometer (Actigraph GT3X+, Actigraph Corp., Pensacola, FL.) and daily sleep diaries along with assessment of daily meal timing/eating patterns. Those with average nightly total sleep time ≥7 h and average daily eating start times within 1 h of awakening were invited to participate in the study. All procedures were approved by the Columbia University Irving Medical Center Institutional Review Board, and all participants provided informed consent form prior to participating in the study.

The main study included two experimental conditions, circadian alignment and circadian misalignment, followed for 6 weeks each. During the circadian alignment condition, participants were instructed to consume their first meal within 1 h of awakening. Two subsequent meals and an evening snack were to be consumed at 5-, 10-, and 11-hours post-awakening, respectively. During the circadian misalignment condition, meals were delayed by 4 hours relative to the aligned condition. For the present analyses, only samples collected during the first 2 weeks of the aligned condition were used. This corresponded to a period during which all meals and snacks were provided to participants based on weight maintenance energy needs, determined using the Mifflin-St. Jeor equation (11). Participants were required to consume all of the foods provided each day and to refrain from consuming other foods or beverages. On days 3-4 and 14-15 of this controlled feeding period, participants underwent cardiometabolic and energy expenditure assessments in a whole-room indirect calorimeter. During these 24-h visits, blood samples were collected every 30 min, from 7 PM to 8:30 AM, via an indwelling catheter connected to an IV line that ran through a port to the outside of the room. Throughout the wake period, participants remained in a semi-recumbent position in dim light, wearing blue-blocking lenses. They were instructed to consume the meals and snack provided at the assigned times, and to turn the lights out at

11:30 PM, to initiate sleep, and were awoken via phone call at 8 AM and instructed to turn the lights back on.

*Serum processing and storage*

Whole blood was collected into 4mL BD vacutainer serum tubes (cat# 367812). Blood remained upright in the tube until clotted, after which it was centrifuged for 20 minutes at 3000 RPM and 4°C. Resultant serum was aliquoted into cryovials and stored at -80°C until GDF15 measurements by ELISA.

#### *4.2 In-house lactate assay*

A subset of plasma lactate levels collected from healthy controls in the *MiSBIE* study (n=54, 76% female) were quantified using lactate assay kits (Abcam, ab65331) following the manufacturer's instructions. Standard curve was prepared and used on each plate; an average standard curve was used for interpolation. Reference samples of known lactate concentrations were used on each plate to monitor batch effect. 16uL of plasma sample was diluted with 184uL of lactate assay buffer to maximize the number of samples within the dynamic range of the assay. Absorbance was gauged at 450nm, and concentrations were interpolated from the trendline equation calculated from the average standard curve.

**Dataset S1 (separate file). Raw GDF15 data from MiSBIE, MaPS, Intensive repeated-measures n=2, Overnight meal alignment study.** Levels of GDF15 were quantified using a high-sensitivity ELISA kit (R&D Systems, DGD150, SGD150) following the manufacturer's instructions, see methods for details. GDF15 measured in pg/mL

**Dataset S2 (separate file). Number of samples by study and sample types, with proportion of missing data.** This table includes information on samples processed in house: MiSBIE, MaPS, Intensive repeated-measures n=2, Overnight meal alignment study. Levels of GDF15 were quantified using a high-sensitivity ELISA kit (R&D Systems, DGD150, SGD150) following the manufacturer's instructions, see methods for details.

**5. Supplemental Information Text References**
