## Supplemental Information Text Figures 1-6 for "The energetic stress cytokine GDF15 is elevated in the context of chronic and acute psychosocial stress"

#### 3. Supplemental Information Text Figures
#### SI Text Figure 1

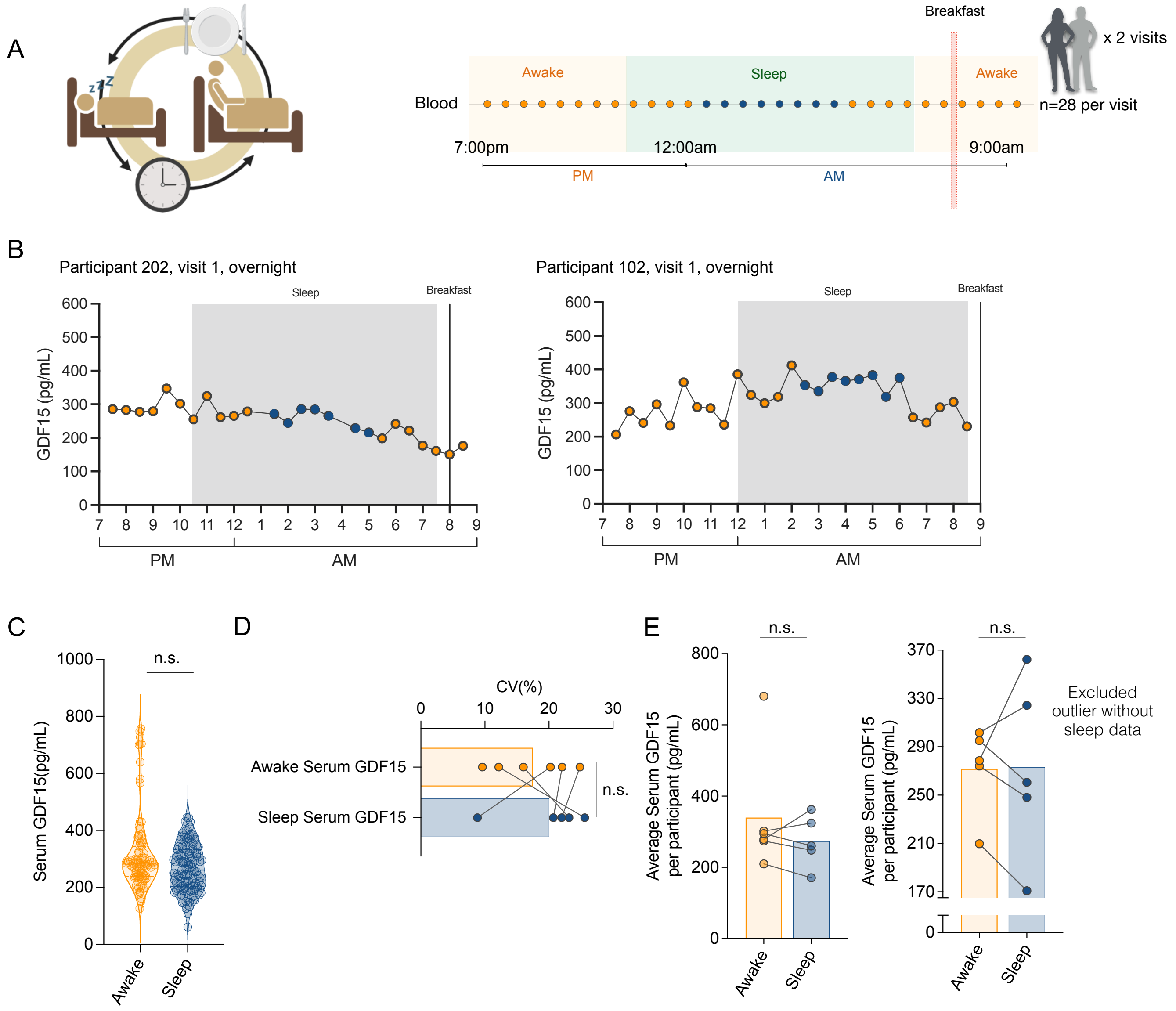


**SI Text Figure 1. Serum GDF15 level does not reliably change between wake to sleep transitions from *Overnight meal alignment study.***
(**A**) *Overnight meal alignment study* experimental design. GDF15 was measured in up to 28 blood samples from n=6 participants during an overnight hospital protocol for two nights. Participants came for two visits where they were sampled once every 30min from 7pm to 9am the next day each visit.(**B**) Full time course shown from a female (*left*) and a male (*right*) participant for one visit. (**C**) Distribution of serum GDF15 measured in healthy adult during sleep and awake time. n=221 observations in 3 females and 3 males, age matched.(**D**) Variability of sleep vs awake serum GDF15 level, each dot reflects one participant's C.V. during sleep and awake time calculated from two visits.(**E**) Average of sleep vs awake levels of GDF15 measured in serum, each dot reflects one participant's C.V. during sleep and awake time calculated from two visits.
P-values from (C) Mann-Whitney t-test and (D,E) Wilcoxon paired t-test, *p<0.05, **p<0.01, ***p<0.001, ****p<0.0001.

### SI Text Figure 2

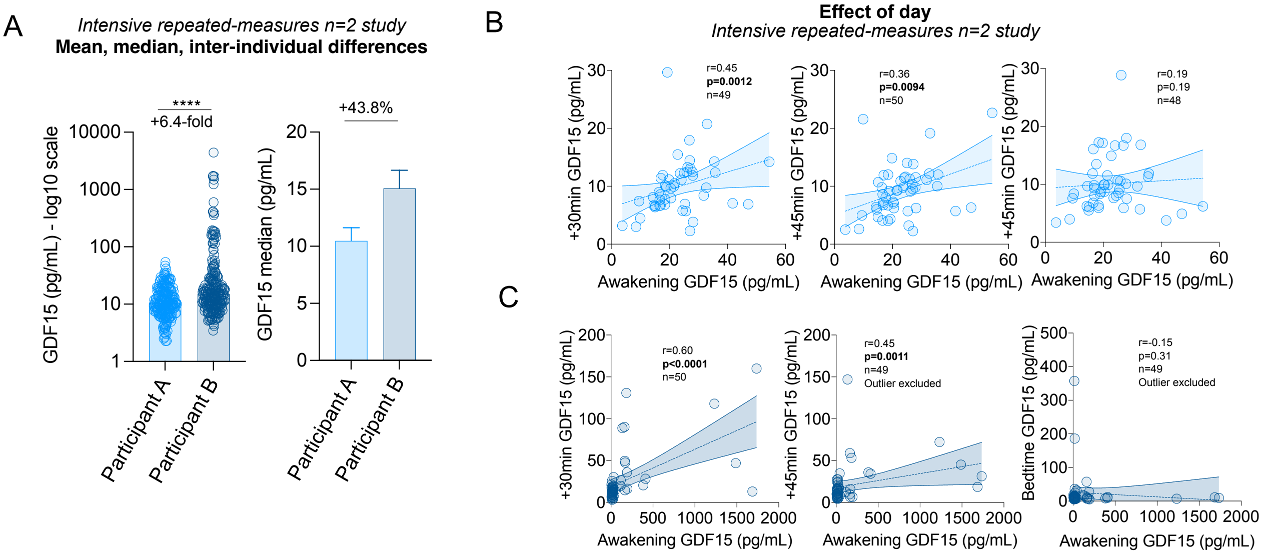


**SI Text Figure 2. Inter-individual differences and effect of day in human saliva GDF15 awakening response.**
(**A**) Inter-individual differences of GDF15 concentrations (*left*, log10 scale; *right*, median on linear scale, data median ± 95% CI) from Participants A and B. (**B**,**C**)Effect of day in Participant A (**B**, 53-day, top) and Participant B (**C**, 60-day, bottom) shown by correlating awakening saliva GDF15 with other measurements in the same day from Intensive repeated-measures n=2 study.
Effect sizes and p-values from (A) Mann-Whitney non-parametric t-test and (B-E) Spearman's rank correlation *p<0.05, **p<0.01, ***p<0.001, ****p<0.0001.

### SI Text Figure 3

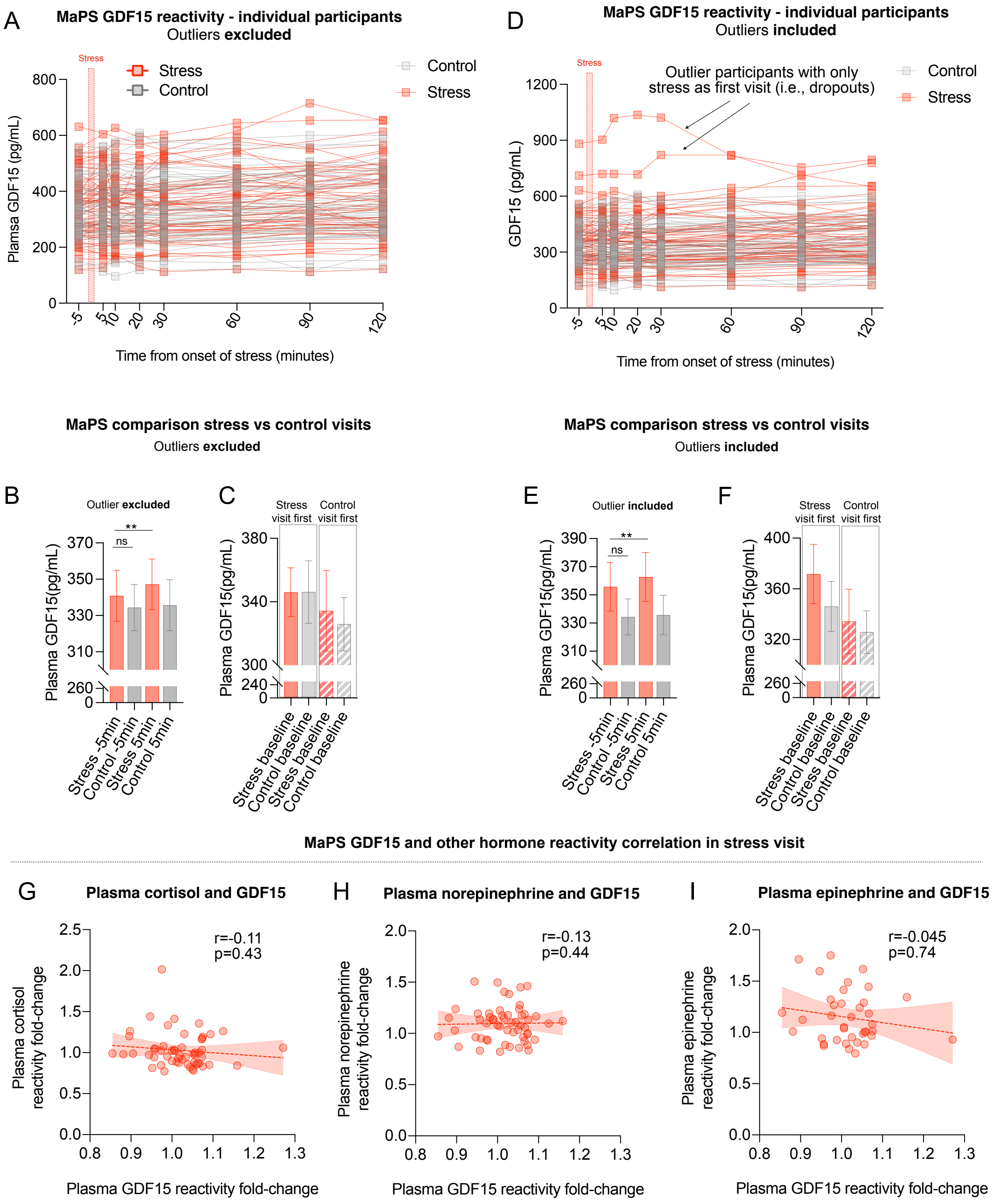


**SI Text Figure 3. Psychological stress acutely induces small-to-moderate changes in plasma GDF15 from MaPS.**
(**A**) Individual plasma GDF15 trajectories for control and stress visits (MaPS, n=68, 49% female, 916 observations; two outliers excluded; see Figure S2D for full data). (**B**) Plasma GDF15 levels at -5min pre-stress (stress visit: n=59, 46% female; control: n=60, 47% female) and 5min post-stress (stress visit: n=58, 45% female; control: n=54, 52% female). (**C**) Baseline (-5 min) plasma GDF15 by visit order (stress first: n=33, 44% female; control first: n=35, 51% female).(**D**) Individual trajectories for all MaPS participants (n=70, 49% female, 916 observations). (**E**) Plasma GDF15 at -5min pre-stress (stress: n=61, 48% female; control: n=60, 47% female) and 5min post-stress (stress: n=60, 47% female; control: n=54, 52% female) with two outliers. (**F**) Baseline (-5 min) plasma GDF15 by visit order with all MaPS participants, including two outliers (stress first: n=35, 46% female; control first: n=35, 51% female). Correlation between plasma GDF15 response and (**G**) cortisol, (**H**) norepinephrine, and (**I**) epinephrine responses during the stress visit (MaPS, n=68, 59% female; two outliers excluded). Plasma GDF15 reactivity was calculated as the fold-change from baseline (-5min) to the group-average peak at 5 minutes. Cortisol stress reactivity was calculated as the fold-change from baseline (-5min) to the peak level at or before 30 minutes post-stress. Norepinephrine and epinephrine stress reactivity was calculated as the fold-change from baseline (-5min) to the peak level at or before 20 minutes post-stress.
(B,C,E,F) Data shown as mean ± SEM, p-values from (B-F) paired non-parametric signed-rank Wilcoxon test and (G-I) Spearman's rank correlation. *p<0.05, **p<0.01, ***p<0.001, ****p<0.0001.

### SI Text Figure 4

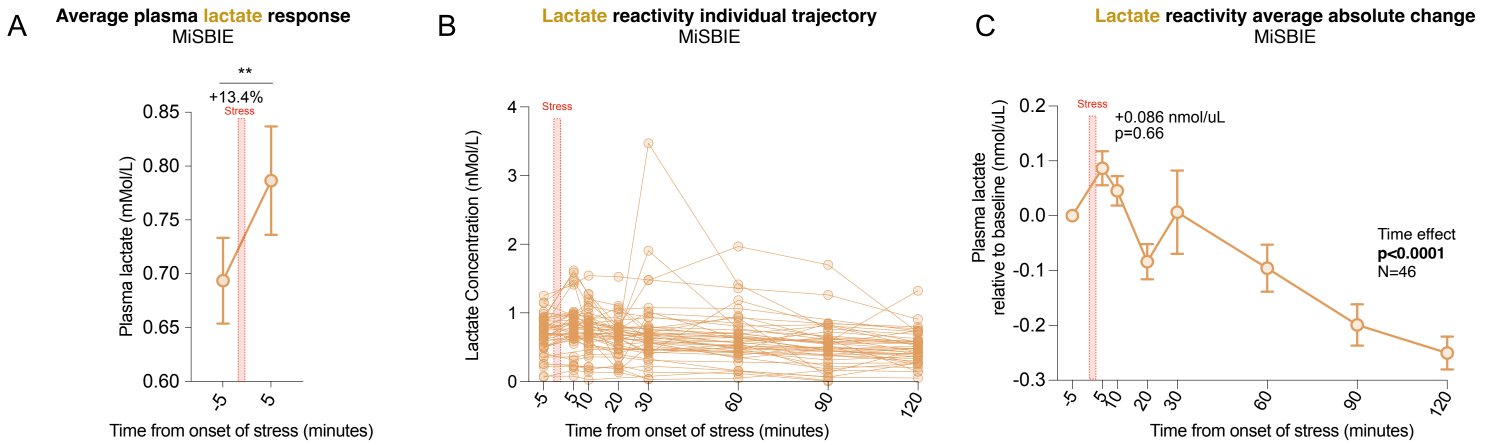


**SI Text Figure 4. Social-evaluative stress acutely increases plasma lactate from a subset of MiSBIE participants**

Plasma lactate levels shown as (**A**) average percent change from baseline (-5 min) and 5min after speech task, (**B**) individual trajectories, (**C**) absolute change from baseline (-5 min). n=46, 72% female, 94 observations.
(A,C) Data shown as mean ± SEM. P-values from (A) Wilcoxon paired t-test and (C) mixed-effect model followed by Tukey multiple comparison test, corrected for multiple testing. *p<0.05, **p<0.01, ***p<0.001, ****p<0.0001.

### SI Text Figure 5

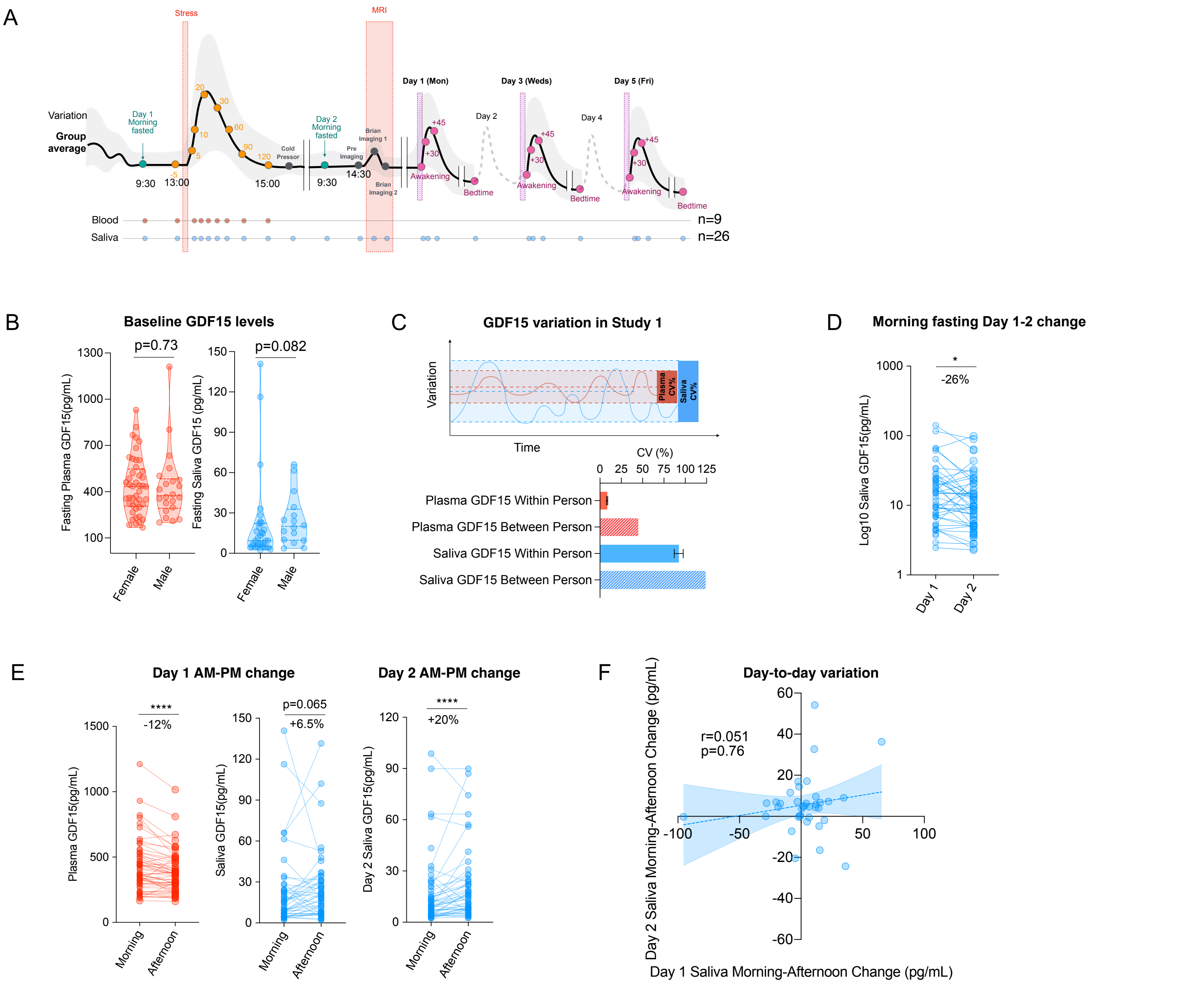


**SI Text Figure 5. Sex comparison and dynamic behavior of GDF15 in human plasma and saliva from MiSBIE**

(**A**) MiSBIE experimental design. GDF15 was measured from human plasma and saliva samples over up to 26 timepoints during a 2-day hospital visit and a 3-day home-based saliva collection protocol (awakening response). (**B**) Sex difference in morning fasting plasma GDF15 and saliva GDF15. Sample sizes for plasma: N=67, 68% female; saliva: n=48, 64% female. Data from morning fasting timepoints. (**C**) Plasma and saliva GDF 15 within person- and between- person variation. Within person coefficient of variation (C.V.) were computed from all timepoints (pictogram above) and between person CV from day 1 morning fasting timepoints. (**D**) Day 1 morning fasting to Day 2 morning fasting saliva levels GDF15. Day 1, n=48 observations, 64% female. Day 2, n=54 observations, 69% females.(**E**) Morning to afternoon levels of GDF15 measured in Day 1 plasma and saliva (*left*) and Day 2 saliva (*right*). Plasma: n=65, 65% female; saliva: n=48, 71% female. (**F**) Correlation between Day 1 and Day 2 saliva GDF15 morning to afternoon change. n=37, 65% female.

Effect sizes and p-values from (B) Mann-Whitney t-test (D,E) Wilcoxon paired t-test, (F) Spearman's rank correlation *p<0.05, **p<0.01, ***p<0.001, ****p<0.0001.

### SI Text Figure 6

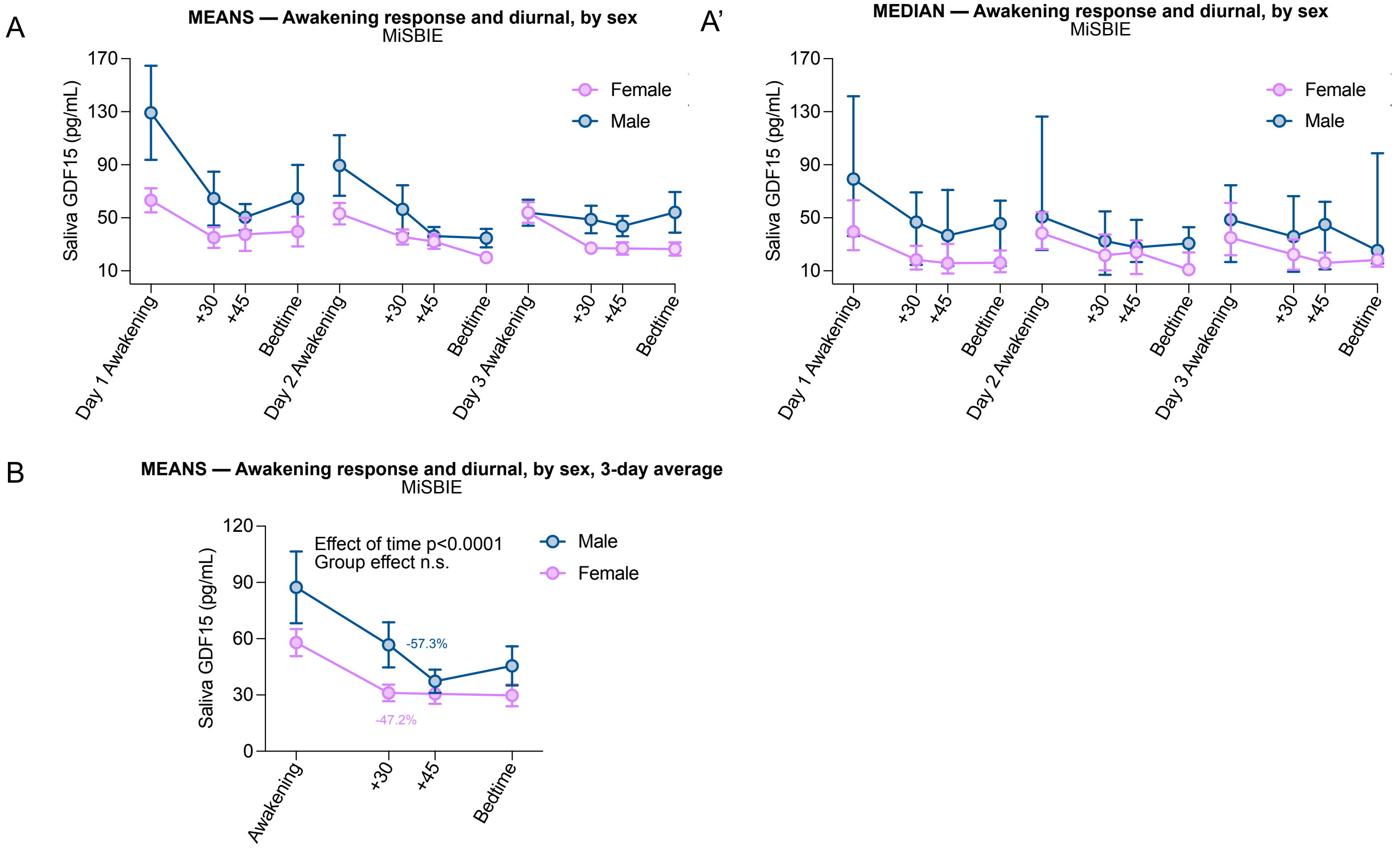


**SI Text Figure 6. Sex comparison of GDF15 awakening response in human saliva.**
(**A,A'**) Sex differences in saliva GDF15 awakening response (female n=48, male n=22) reported as average (A) and median (A') from MiSBIE. (**B**)Sex differences in saliva GDF15 3-day average awakening response (female n=48, male n=22) from MiSBIE. P-values for group effect from two-way ANOVA followed by Tukey multiple comparison test for effect of time, corrected for multiple testing.
